# Structural context of homomeric interactions in the lg domain of the MPZ (P0) myelin adhesion protein and relation to Charcot-Marie-Tooth disease phenotype variants

**DOI:** 10.1101/2023.03.18.533291

**Authors:** Christopher P. Ptak, Tabitha A. Peterson, Jesse B. Hopkins, Christopher A. Ahern, Michael E. Shy, Robert C. Piper

## Abstract

Mutations in Myelin Protein Zero (MPZ) account for 5% of Charcot-Marie-Tooth cases and can cause demyelinating or axonal phenotypes, reflecting the diverse roles of MPZ in Schwann cells. MPZ holds the apposing membranes of the myelin sheath together, with the adhesion role fulfilled by the extracellular lmmunoglobulin-like domain (lg^MPZ^), which can oligomerize. Current knowledge for how the lg^MPZ^ might form oligomeric assemblies involving 3 weakly-interacting interfaces has been extrapolated from a protein crystal structure in which individual rat lg^MPZ^ subunits are packed together under artificial conditions. These interfaces include one that organizes the lg^MPZ^ into tetramers, a ‘dimer’ interface that could link tetramers together, and a third hydrophobic interface that could mediate binding to lipid bilayers or the same hydrophobic surface on another lg^MPZ^ domain. There are at present no data confirming whether the proposed lg^MPZ^ interfaces actually mediate oligomerization in solution, whether they are required for the adhesion activity of MPZ, whether they are important for myelination, or whether their loss results in disease. We performed NMR and SAXS analysis of wild-type lg^MPZ^ as well as mutant forms with amino-acid substitutions designed to interrupt its presumptive oligomerization interfaces. Here, we confirm the interface that mediates lg^MPZ^ tetramerization, but find that dimerization is mediated by a distinct interface that has yet to be identified. We next correlated CMT phenotypes to subregions within lg^MPZ^ tetramers. Axonal late-onset disease phenotypes (CMT2l/J) map to surface residues of lg^MPZ^ proximal to the transmembrane domain. Early-onset demyelinating disease phenotypes (CMT1B/Dejerine-Sottas syndrome) map to two groups: one is described by variants that disrupt the stability of the lg-fold itself and are largely located within the core of the lg domain; whereas another describes a surface on the distal outer surface of lg^MPZ^ tetramers. Computational docking studies predict that this latter disease-relevant subregion may mediate dimerization of lg^MPZ^ tetramers.

## INTRODUCTION

Charcot Marie Tooth (CMT) disease refers to heritable peripheral neuropathies, which are the most common genetic neuromuscular diseases, affecting 1:2500 individuals (Skre, 1974). Autosomal dominant (AD) inheritance is the most common, followed by X-linked and autosomal recessive (AR) forms. Most forms of CMT are demyelinating while one-third are primary axonal disorders (Fridman et al., 2015; Murphy et al., 2012). Mutations in Myelin Protein Zero (MPZ) account for 5% of CMT cases and can cause demyelinating (CMT1B) or axonal phenotypes (CMT2l/J), likely reflecting multiple molecular functions of MPZ in Schwann cells (Fridman et al., 2015; Hayasaka, Himoro, Sato, et al., 1993).

MPZ comprises ∼50% of peripheral nervous system (PNS) myelin proteins and is necessary for normal myelin structure and function (Eylar et al., 1979; Greenfield et al., 1973). MPZ is a single-pass transmembrane protein consisting of a single immunoglobulin-like extracellular domain (Lemke & Axel, 1985; Uyemura et al., 1995) and cytosolic tail (*Supplemental Figure 1*); that can be post-translationally modified by N-linked glycosylation, sulfation, palmitoylation, and phosphorylation (D’Urso et al., 1990; Eichberg & lyer, 1996). One function of MPZ is as an adhesion protein, holding together adjacent wraps of myelin membrane, which is thought to be mediated in part through homotypic interactions of its extracellular lg domain (Filbin et al., 1990; Giese et al., 1992; Xu et al., 2001). The cytosolic tail of MPZ has a signaling role, promoting myelin compaction of the cytosolic region between myelin wraps to form the major dense line (Filbin et al., 1999; Mandich et al., 1999; Xu et al., 2001).

Exactly how the lg domain of MPZ (lg^MPZ^) mediates adhesion of apposing membranes remains to be fully determined, and is important for understanding how myelin is constructed and how mutations in the lg^MPZ^ cause disease. Available models hypothesize that homotypic oligomerization of the lg^MPZ^ mediates membrane adherence (Filbin et al., 1990; Giese et al., 1992; Xu et al., 2001). Yet, biochemically verified interactions that support *cis* and *trans* coupling of MPZ via its extracellular lg domain are unknown (Filbin et al., 1999; Mandich et al., 1999; Xu et al., 2001). Our current understanding for how the lg^MPZ^ forms oligomeric assemblies has been extrapolated from a protein crystal structure in which individual rat lg^MPZ^ subunits were packed together under artificial conditions (Shapiro et al., 1996). This arrangement indicated two protein:protein interfaces: an asymmetric interface that drives the assembly of lg^MPZ^ tetramers that would bundle 4 MPZ proteins embedded in the same (*cis*) membrane; and a symmetrical dimeric interface involving MPZ in apposing (*trans*) membranes. While this model is further rationalized by its fit within the intraperiod line of myelin that spans the gap between adhered plasma membrane sheaths of myelin, crystal packing alone is an insufficient measure of how oligomerization occurs in solution. Moreover, there are additional data suggesting that this model may not account for how normally lg^MPZ^ oligomerizes. For instance, another crystal structure of the human lg^MPZ^ has a different crystal packing suggesting that the potential binding interfaces in the rat lg^MPZ^ crystal lattice may not be dominant drivers of oligomer assembly (Liu et al., 2012). Recently, low resolution cryo-EM data from SDS-solubilized MPZ isolated from bovine nerve led authors to propose a different configuration for how lg^MPZ^ mediates adhesion (Raasakka et al., 2019), namely that *cis*-tetramers might not form but rather *trans*-dimers interacting as a zipper to drive membrane adhesion. Finally, while CMT-causing disease variants within the lg^MPZ^ are near the interfaces identified by crystal packing (Grandis et al., 2004; Raasakka & Kursula, 2020; Shy, 2005; Shy, 2006), there are no disease-causing variants that are known to precisely alter the surface residues of these interfaces. Together, these latter observations present two non-exclusive possibilities: 1) that the interfaces thought to control lg^MPZ^ oligomerization are incorrect and 2) that the lg^MPZ^ oligomerization is not strictly required for myelin function or that its interruption may not lead to a dominant disease phenotype that would present clinically. At least one pathogenesis pathway that causes demyelinating CMT has been identified that involves accumulation of misfolded MPZ in the ER, evoking ER-stress and activating the unfolded protein response (UPR) (Pennuto et al., 2008; Saporta et al., 2012). However, not all disease-causing variants retain MPZ in the ER and/or cause ER stress suggesting there must be other pathogenic pathways that possibly involve disruption of protein-protein interactions of lg^MPZ^ (Bai et al., 2018).

To lend insight into the role of MPZ oligomerization for normal myelin formation, we performed solution binding studies that examined whether lg^MPZ^ oligomerization is mediated through the interfaces identified by previous crystal packing data. We performed NMR and SAXS analysis of wild-type lg^MPZ^ as well as mutant forms with amino-acid substitutions designed to interrupt its presumptive oligomerization interfaces. Our data confirm that the lg^MPZ^ domain does form tetramers in solution, oriented in *cis* with respect to the membrane. We also confirm that these tetramers dimerize, however their mode of dimerization is inconsistent with current models and appears mediated by an interface that remains to be identified. To lend insight into the role of MPZ oligomerization for normal myelin formation and to provide a structural correlate for pathogenesis of CMT, we performed solution binding studies that examined whether lg^MPZ^ oligomerization is mediated through the interfaces identified by previous crystal packing data and then analyzed the location of dominant disease-causing amino-acid substitutions within the lg^MPZ^ domain. To better understand lg^MPZ^ function and provide insight into whether distinct molecular mechanisms might drive CMT pathogenesis, we mapped disease causing patient variants onto the individual lg^MPZ^ structure as well as that of its *cis*-tetrameric structure. Here, we identify 3 spatially distinct regions. First, a subset of variants that cause severe early-onset demyelinating CMT1B and DSS (Dejerine-Sottas Syndrome) map to the core of the lg^MPZ^ domain. Computationally, these variants are predicted to de-stabilize the integrity of the lg fold, consistent with previous studies showing that these variants when tested, evoked the UPR and cause MPZ to accumulate in the ER. A second set of variants map to the distal side of MPZ tetramers, potentially mediating interactions with other proteins such as MPZ from an apposing membrane. A third subregion mapping the proximal surface of the lg^MPZ^ tetramers correlates with late-onset (CMT2l/J) in which myelin is mostly normal but axons degenerate (McCray & Scherer, 2021).

Finally, we integrate a computational evaluation of disease-causing variants on the lg^MPZ^ surface with alternative hypothetical oligomerization interfaces and discuss the possibilities for how MPZ mediates adhesion of myelin layers and how its inability to do so might mediate different forms of CMT.

## MATERIALS AND METHODS

### Plasmids

Plasmids used for protein expression are listed in *Supplemental Table 1*. Gibson assembly cloning (NEB BioLabs, lpswich, MA) was used to make plasmids. pPL7238, a low-copy pET-based plasmid used for lg^MPZ^ expression in *E.coli* contains a T7 RNA polymerase promoter followed by coding regions for Maltose Binding Protein (MBP) (Baneyx & Mujacic, 2004) containing an N-terminal bacterial secretion tag; a tabacco-etch virus protease site; an AASM linker, and residues l30-R153 of human MPZ (NP_000521.2) as encoded in a synthetic DNA fragment (lDT, Coralville, lA) recoded to the *E. coli* codon bias. The MPZ-eGFP-HA expression plasmid pPL7430 was made using a human codon optimized synthetic DNA fragment encoding M1-K248 of MPZ (NP_000521.2), a TGSGS linker, eGFP and HA epitope tag cloned into pcDNA3.1 downstream of the CMV promoter. All point mutations of plasmids were generated by Quick Change PCR mutagenesis using Pfu Ultra polymerase (Agilent Technologies, Santa Clara, CA).

### Protein expression and purification

MBP-lg^MPZ^ fusion proteins were expressed in Origami 2(DE3) competent *E. Coli* (Novagen, Reno, NV) by induction with 0.5 mM lsopropyl β-d-1-thiogalactopyranoside (lPTG) at 26.5 °C for 18 hr in kanamycin-supplemented LB media or M9-based minimal media with ^15^NH_4_Cl, trace elements, CaCl_2_, MgSO_4_, Thiamine, and 0.3% glucose. The periplasmic fraction containing the expressed fusion protein was isolated by osmotic shock (Shapiro et al., 1996) where fresh cell pellets from 4 L of culture were solubilized in 1.6 L of 20% sucrose, 30 mM TrisCl (pH 8), followed by addition of 1 mM EDTA and stirring for 8 min at 23 °C. The cell suspension was centrifuged at 6000 rpm for 30 min at 4 °C. After discarding the supernatant, the pellet was resuspended in 1.6 L of ice-cold 5 mM MgSO_4_ and stirred for 10 min at 4 °C. The suspension was centrifuged at 5000 rpm for 20 min at 4 °C and the resulting supernatant containing the periplasmic isolate was placed on ice. Protease inhibitors (Complete EDTA-free, Roche, Basel, CH) and 20 mM TrisCl (pH 7.6) were added to the periplasmic isolate, which was sterile filtered and concentrated on DEAE-Sepharose fast flow resin (Amersham Biosciences, Buckinghamshire, UK). The periplasmic isolate was eluted with column buffer (150 or 200 mM NaCl, 1 mM EDTA, 1 mM NaAzide, 20 mM TrisCl (pH 7.6)) and applied to Amylose resin (New England BioLabs, lpswich, MA). The purified MBP-lg^MPZ^ fusion protein was eluted with column buffer containing 10 mM maltose and stored at 4 °C. Purified MBP-lg^MPZ^ fusion protein was diluted 3-fold with 20 mM TrisCl (pH 7.6), and incubated with TEV protease for 16 hr at 23 °C. The cleaved proteins were separated by a 16/600 Superdex75 gel filtration column (GE Healthcare, Chicago, lL), followed by depleting the peak of lg^MPZ^ peak or residual MBP by by applying to amylose resin, and finally a second HiLoad 16/600 Superdex75 gel filtration column, each run in 50 mM NaCl, 1 mM EDTA, 1 mM NaAzide, 20 mM TrisCl (pH 7.6).

### NMR Spectroscopy

For NMR experiments, ^15^N-labeled lg^MPZ^ proteins were exchanged into NMR buffer (25 mM NaCl, 0.5 mM EDTA, 0.5 mM NaAzide, 10 mM TrisCl (pH 6.9), 10% D_2_O) and concentrated to 160-190 µM. Final protein samples were measured at 1x, 0.5x, and 0.25x concentrations diluted with NMR buffer. Spectra were collected at 20 °C on a Bruker AVANCE NEO 600 MHz NMR spectrometer with a gradient cryoprobe. ^15^N/^1^H HSQC spectra were processed using NMRPipe (Delaglio et al., 1995) and analyzed using POKY (Lee et al., 2021). Longitudinal (T_1_) and transverse (T_2_) ^1^H-detected 1D ^15^N relaxation spectra were acquired on protein samples and relaxation curve-fitting analysis was performed with Topspin 4.0 (Bruker, Billerica, MA). For comparison with experimentally determined T_1_/T_2_ ratios, a theoretical T_1_/T_2_ ratio for the lg^MPZ^ monomer was determined using HydroNMR (de la Torre et al., 2000) on NMRBox (Maciejewski et al., 2017).

### SAXS analysis

For SAXS experiments, lg^MPZ^ proteins were exchanged into SAXS buffer (50 mM NaCl, 1 mM EDTA, 20 mM TrisCl (pH 7.6). lg^MPZ^ wild-type and W53A, R74A, D75R mutant were concentrated to 105 and 200 µM, respectively, and were subsequently diluted to the appropriate concentration (50-200 µM) with SAXS buffer. The final 50 µl samples were centrifuged for 5 min at 13,000 rpm. SAXS was performed at BioCAT (beamline 18lD at the Advanced Photon Source, Chicago) with in-line sample loading that was run at 0.3-0.4 ml/min on an AKTA Pure FPLC (GE) without column separation. The flow cell consists of a 1.0 mm lD quartz capillary with ∼20 µm walls. A coflowing buffer sheath was used to separate sample from the capillary walls and to help prevent radiation damage (Kirby et al., 2016). Scattering intensity was recorded using an Eiger2 XE 9M (Dectris) detector placed 3.65m from the sample giving us access to a q-range of 0.003 A^-1^ to 0.34 A^-1^ (for wild-type) or 0.003 A^-1^ to 0.42 A^-1^ (for the W53A, R74A, D75R mutant). 0.25 s exposures were acquired every 0.5 s during elution.

Data were reduced using BioXTAS RAW 2.1.1 (Hopkins et al., 2017). l(q) vs q scattering curves were created from exposures selected from the sample peak. Matching buffer blanks for background subtraction were obtained from averaged pre-sample exposures. The fraction of oligomers was determined from experimental SAXS data using the OLlGOMER program from the ATSAS suite (Petoukhov et al., 2012). Homo-oligomeric arrangements of lg^MPZ^ were identified from the expanded unit cell of the lg^MPZ^ crystal structure (PDB lD: 1NEU)(Shapiro et al., 1996) generated in UCSF Chimera (Pettersen et al., 2004). To obtain theoretical form factors, structural predictions for the final proteins from our expressed human lg^MPZ^ constructs were generated by a ColabFold (Mirdita et al., 2022) search of MMseqs2 (Steinegger & Soding, 2017) with AlphaFold2 (Jumper et al., 2021). Oligomeric arrangements were generated by superimposition of ColabFold lg^MPZ^ structures onto crystal structure-derived lg^MPZ^ oligomers (Meng et al., 2006).

### Fluorescence microscopy

HEK293 cells were cultured in Dulbecco’s modified Eagle’s medium (Gibco) supplemented with 10% fetal bovine Serum (Gibco) and 50 μg /mL penicillin/streptomycin at 37 °C with 5% CO_2_. Cells were seeded on glass bottom dishes. Plasmid transfections were carried out in the absence of penicillin/streptomycin using Lipofectamine LTX Reagent with Plus Reagent (lnvitrogen, Cat# A12621). 24 hr post transfection live cell micrographs were captured using a Leica SP8 confocal microscope equipped with a 488 nm laser for excitation and an emission collection window of 520-540 nm.

### Structural Analysis

Crystal Packing of the lg^MPZ^ domain within 1NEU was examined using PlSA (Krissinel & Henrick, 2007). Predictions on the effects of mutations on the interfaces predicted by the 1NEU crystal structure were made using SSlPe (Huang et al., 2020). Calculating values for accessible surface area for each amino acid in the lg^MPZ^ of 3OAl was done using UCSF Chimera (Pettersen et al., 2004). Distance measurements and clash scores were also calculated using UCSF Chimera. Prediction of changes in the stability of the lg^MPZ^ of 3OAl with different missense mutations was done using DeepDDG (Cao et al., 2019). Generation and refinement of alternative lg^MPZ^ dimer interfaces were done using ClusPro2 (Kozakov et al., 2017).

## RESULTS

### Oligomerization interfaces for lg^MPZ^

The lg^MPZ^ oligomerizes and this property is thought to mediate adhesion of myelin wraps (Lemke & Axel, 1985). The interfaces hypothesized to mediate MPZ oligomerization have been extrapolated from a crystal structure of the rat lg^MPZ^ wherein individual lg^MPZ^ units form interfaces within the crystal lattice (Shapiro et al., 1996). The arrangement of these 3 interfaces and how they have been thought to bridge the intraperiod line are schematically shown in Figure 1. One heterotypic interface (lnterface A) would mediate formation of lg^MPZ^ tetramers in a parallel ‘*cis’* orientation emanating from the same membrane. A homotypic dimer interface (lnterface B) would link two apposing lg^MPZ^ in antiparallel ‘*trans’* configuration that spans the ∼45 A intraperiod line. ln the model by Shapiro et al., this ‘dimer’ interface connects 2 MPZ tetramers each emanating from apposing membranes in the myelin wrap (Shapiro et al., 1996). A third interface (lnterface C) in the rat crystal lattice highlights a hydrophobic patch on the distal surface of the lg domain, containing tryptophan residues that could mediate hydrophobic interactions with the apposing lipid bilayer. An alternate model (Figure 1C) lacks MPZ *cis* tetramers and instead proposes that the dimer interface mediates *trans*-paired interaction of two MPZ proteins that work with other MPZ *trans* pairs to zipper myelin wraps together (Raasakka & Kursula, 2020; Raasakka et al., 2019).

**Figure 1.**
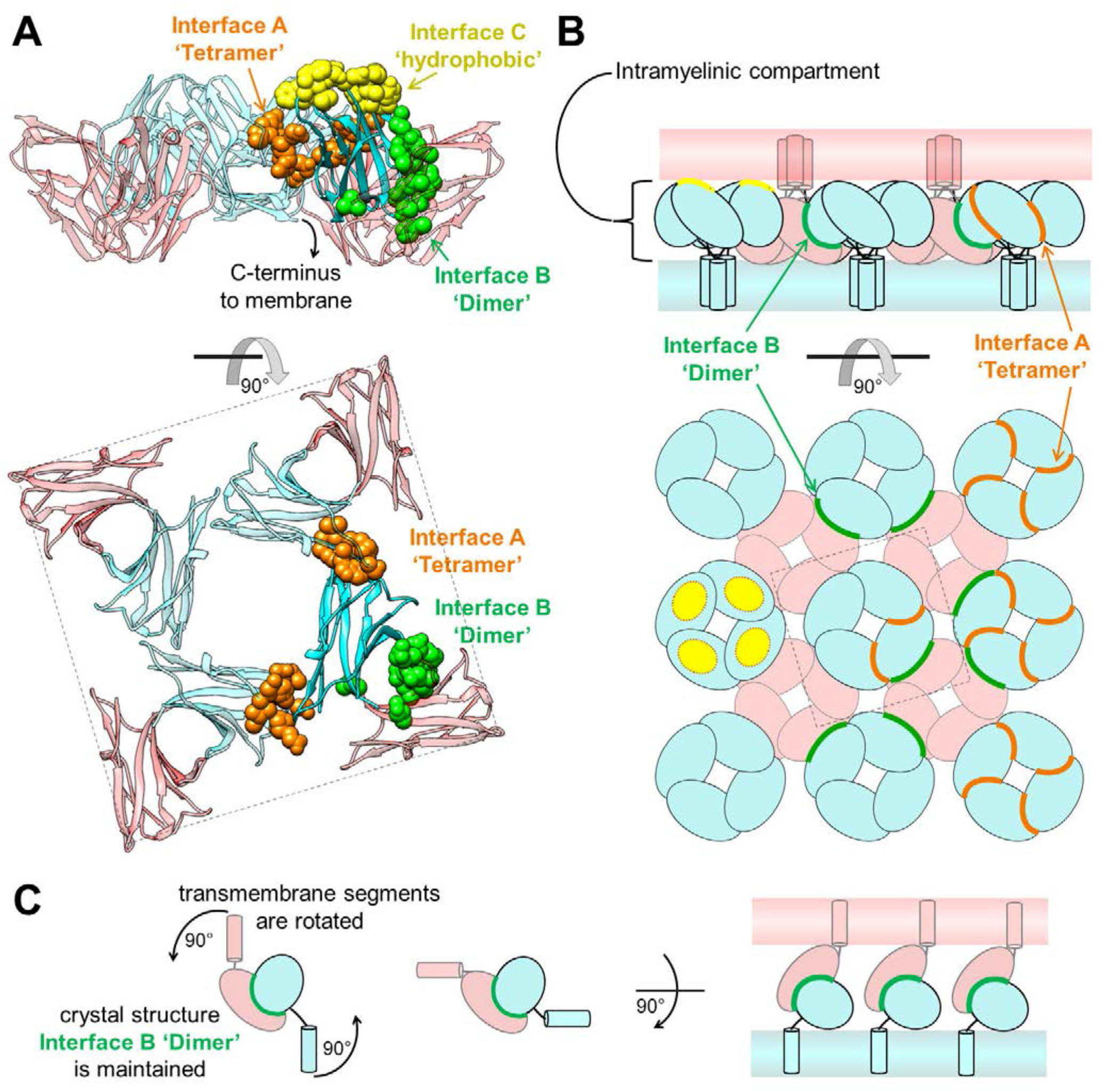
Current models for structural organization of lg^MPZ^ functional assembly in myelin. **(A)** Intermolecular packing in the rat lg^MPZ^ crystal structure (PDBid: 1NEU) (Shapiro, et al., 1996). lg^MPZ^ subunits organize into tetramers (cyan) with the C-terminus on the same side in a *cis* configuration. Each subunit in the tetramer is also dimerized to another lg^MPZ^ subunit from an apposing membrane in a *trans* configuration (pink). Surface residues mediating the *cis*-tetramer (Interface A; orange) and *trans*-dimer (Interface B; green) interactions are displayed. Shown in yellow is another proposed interface made of hydrophobic residues that may associate directly with the lipid bilayer. **(B)** The oligomeric assembly model in A, schematized and extended in the context of the intraperiod line (top). Tetramers in one membrane, held together by the orange tetramer Interface A, assemble with other tetramers of the apposing membrane via the green dimer Interface B. **(C)** An alternate proposed model, lacking a *cis*-tetramer lg^MPZ^ where myelin wraps are held together by two apposing lg^MPZ^ domains interacting through Interface B.

Although these are plausible models, there are few data corroborating how lg^MPZ^ oligomerizes. Moreover, it is not clear whether lg^MPZ^ oligomerization is the driver for adhesion nor whether any of the hypothetical interfaces described in Figure 1 are critical for adhesion. Of note, the interfaces in the crystal structure are predicted to be crystallization artifacts by PlSA. Further, it is unclear whether disruption of oligomeric interfaces would cause dominant negative demyelinating and/or axonal CMT.

### Empirical testing of lg^MPZ^ oligomerization interfaces

To investigate how lg^MPZ^ oligomerizes in solution, we purified recombinant human lg^MPZ^ secreted from *E. coli* and analyzed it at various concentrations by small X-ray scattering (SAXS) analysis and by NMR. ln addition, we also analyzed mutant lg^MPZ^ containing amino acid substitutions chosen to disrupt lnterface A (tetramer) alone, or in combination with disruption of lnterface B (dimer). Figure 2A shows the residues involved in lnterface A and B. lnterface A centers on W53 in one lg^MPZ^ subunit, which is coordinated by H115, E97, and N116 on another lg^MPZ^ subunit. lnterface B defines a symmetric dimer held together by R74, D75 on one monomer engaging S78 and H81 on the other. Using a prediction algorithm, (Huang et al., 2020; Pahari et al., 2020; Rodrigues et al., 2019) we queried which different amino acid substitutions we could engineer that would cause the greatest change in binding energy at these interfaces (Figure 2A and *Supplemental Table 2*). This analysis identified the W53A mutation to disrupt lnterface A (ΔA), and R74A, D75R mutations to disrupt lnterface B (βB). None of these mutations have been reported in CMT patients, thus serving as the basis of an unbiased structure/function analysis.

**Figure 2.**
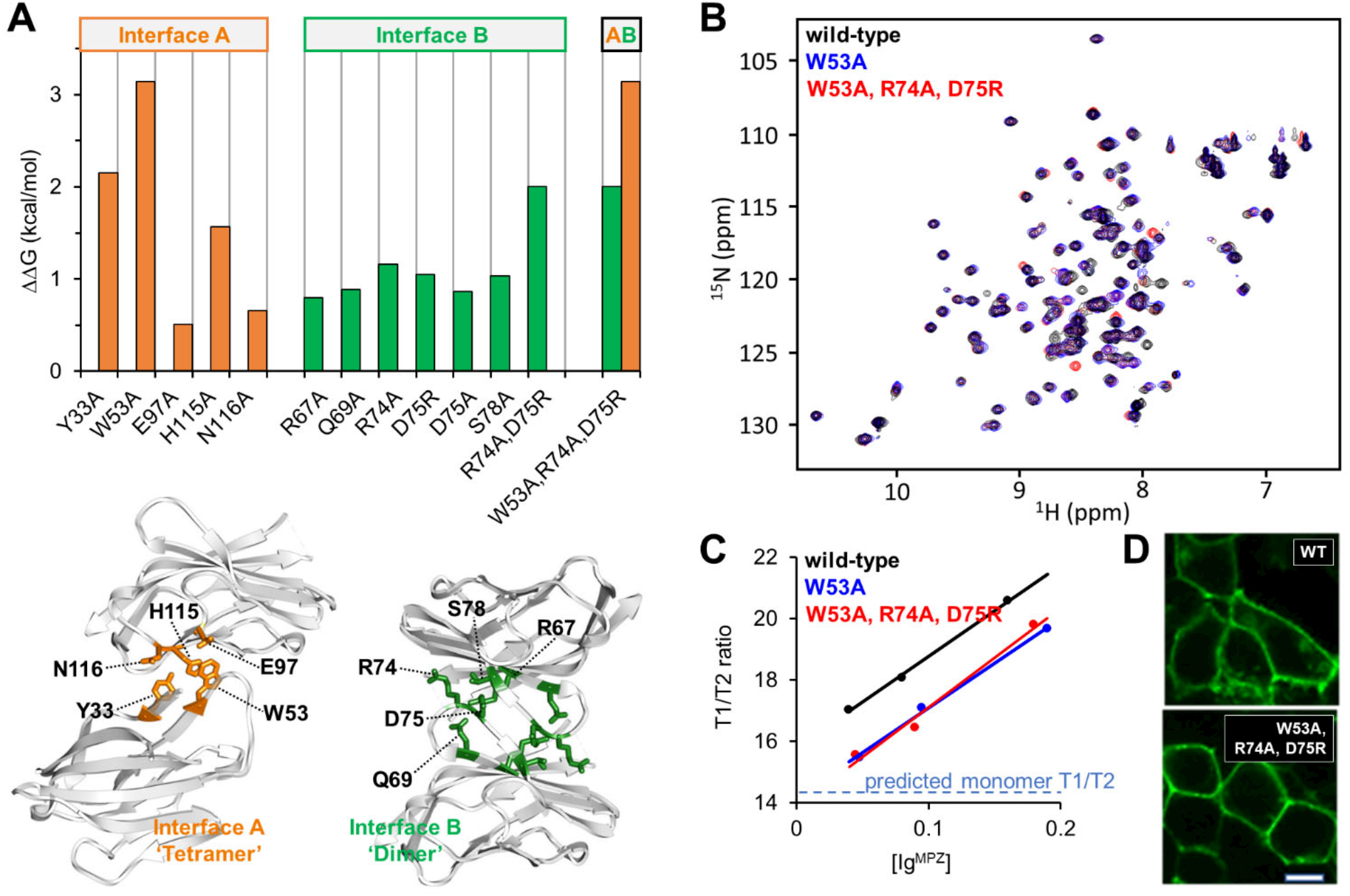
Engineered mutations targeting crystal structure dimer interface studied by solution nuclear magnetic resonance. Targeted mutagenesis coupled with NMR analysis of lg^MPZ^ confirms location of Interface A but not Interface B. **(A)** (Top) To rationally design mutations that specifically distrupt protein interaction interfaces (bar labels above denote intended interface target), we used a computational approach to predict changes in binding energy for indicated amino acid substitutions within lg^MPZ^ for the tetramer Interface A (orange) and dimer Interface B (green). Note that all mutations targeting Interface A, including W53A, only have predicted effects on Interface A and not Interface B. Likewise all mutations targeting Interface B, including the double residue subsitution R74A, D75R, only have predicted effects on Interface B. The W53A, R74A, D75R combination mutant is predicted to comprimise both Interface A and B (data are graphed from calculations shown in *Supplemental Table 2*) (Bottom) Structural diagrams of lg^MPZ^ dimers bound via Interface A (left, orange) or Interface B (right, green). Interface mutants expected to cause the largest oligomeric disruptions were the top candidates to evaluate experimentally. **(B)** Overlay of ^15^N/^1^H HSQC NMR spectra of ^15^N-labeled wild-type lg^MPZ^ (black), the W53A mutant intended to disrupt the Tetrameric Interface A (blue) lg^MPZ^ 11A, and the combined W53A, R74A, D75R triple mutant targeting both Interface A and the dimer Interface B (red) lg^MPZ^ 11A11B. Most ^15^N/^1^H HSQC peaks of the 3 diferent proteins overlap showing that they share high degree of structural similarity and stability (individual spectrum displayed in *Supplemental Figure 2*) **(C)** NMR analysis of molecular tumbling. The ratio of T1 and T2 relaxation times for ^15^N-labeled wild-type lg^MPZ^, lg^MPZ^ 11A, and lg^MPZ^ 11A11B were measured and plotted as a function of protein concentration. An increase in T1/T2 ratio corresponds to an increase in molecular size related to oligomerization. For comparison, the predicted T1/T2 ratio of monomer only (blue hashed line) is also plotted. **(D)** Localization of C-terminally GFP-tagged wild-type MPZ (WT) or mutant MPZ 11A11B (W53A, R74A, D75R) expressed in HEK293 cells via transient transfection. MPZ 11A11B trafficked to the cell surface similar to wild-type MPZ showing both proteins were not retained in the endoplasmic reticulum (ER) and passed ER quality control.

We then analyzed ^15^N-labeled wild-type lg^MPZ^, lg^MPZ^ΔA (W53A) and lg^MPZ^ΔAΔB (W53A, R74A, D75R) domains in NMR ^15^N/^1^H HSQC experiments (Figure 2B and C). All produced similar well dispersed spectra of backbone amides indicating all three lg^MPZ^ domains were well-folded as expected since the mutations chosen targeted surface exposed residues that are not used to support the integrity of the lg fold itself. ln the context of full-length MPZ with a C-terminal tagged GFP, both wild-type and the ΔAΔB mutant were found primarily at the cell surface of transiently transfected HEK293 cells, indicating each had folded properly in cells and were not retained by the ER quality control machinery (Figure 2D). We next measured the average T1 (longitudinal) and T2 (transverse) relaxation times, the ratio of which is proportional to the size of lg^MPZ^ complexes (Figure 2C). T1/T2 measurements were performed for a series of lg^MPZ^ concentrations. The T1/T2 data for wild-type lg^MPZ^ showed a poor fit to monomer alone as predicted by HydroNMR (Garcia de la Torre et al., 2000), and instead showed an increasing T1/T2 ratio with increasing protein concentrations indicating that oligomers assemble in the affinity range of ∼100 µM. ln contrast, both the lg^MPZ^ΔA and lg^MPZ^ΔAΔB mutants had a lower T1/T2 ratio across the same concentration range indicating that the extent of oligomerization was different from that of wild-type lg^MPZ^. However, the T1/T2 ratio of lg^MPZ^ΔA and lg^MPZ^ΔAΔB still increased in a concentration-dependent manner and were indistinguishable from one another, suggesting that oligomerization still occurred. One likely possibility is that there are indeed more than one interface driving oligomerization and that the W53A mutation disrupted one of these (lnterface A), but that the other(s) interface was unaffected by mutating residues that mediate interaction through the hypothetical dimer interface (lnterface B).

While T1/T2 data can differentiate between fully monomeric lg^MPZ^ vs mixtures containing a proportion of lg^MPZ^ oligomers, it lacks the resolution to differentiate between types of oligomers (molecules > 25 kDa, e.g., dimers, tetramers). SAXS, a complementary solution based method with increasing resolution at higher molecular weights, was used to discern the oligomeric status of wild-type lg^MPZ^ at different concentrations (Figure 3A, *Supplemental Figure 3*, and *Supplemental Table 3*). Experimental SAXS curves generated from lg^MPZ^ samples were fit to a mixture of curves computed from theoretical protein models of their predicted homo-oligomeric components. The proportions of each component curve were used to deduce the proportion of oligomers in the solution. The set of component oligomers in the fitting model were optimized to obtain computed SAXS curves with the best fits to empirical SAXS data. Using OLlGOMER (Konarev et al., 2003), our empirical SAXS data fit poorly to theoretical predictions containing only monomeric lg^MPZ^ (x^2^>25). ln contrast, a theoretical curve generated from 5 oligomeric components had a very good fit (x^2^=0.99). The minimal set of lg^MPZ^ oligomers to obtain a reasonable fit (x^2^<1.5) required the inclusion of both the monomer and tetramer. At the higher concentration, the equilibrium shifts away from monomers and more to dimers, tetramers, 8-mers, and 16-mers (Figure 3C and *Supplemental Table 4*).

**Figure 3.**
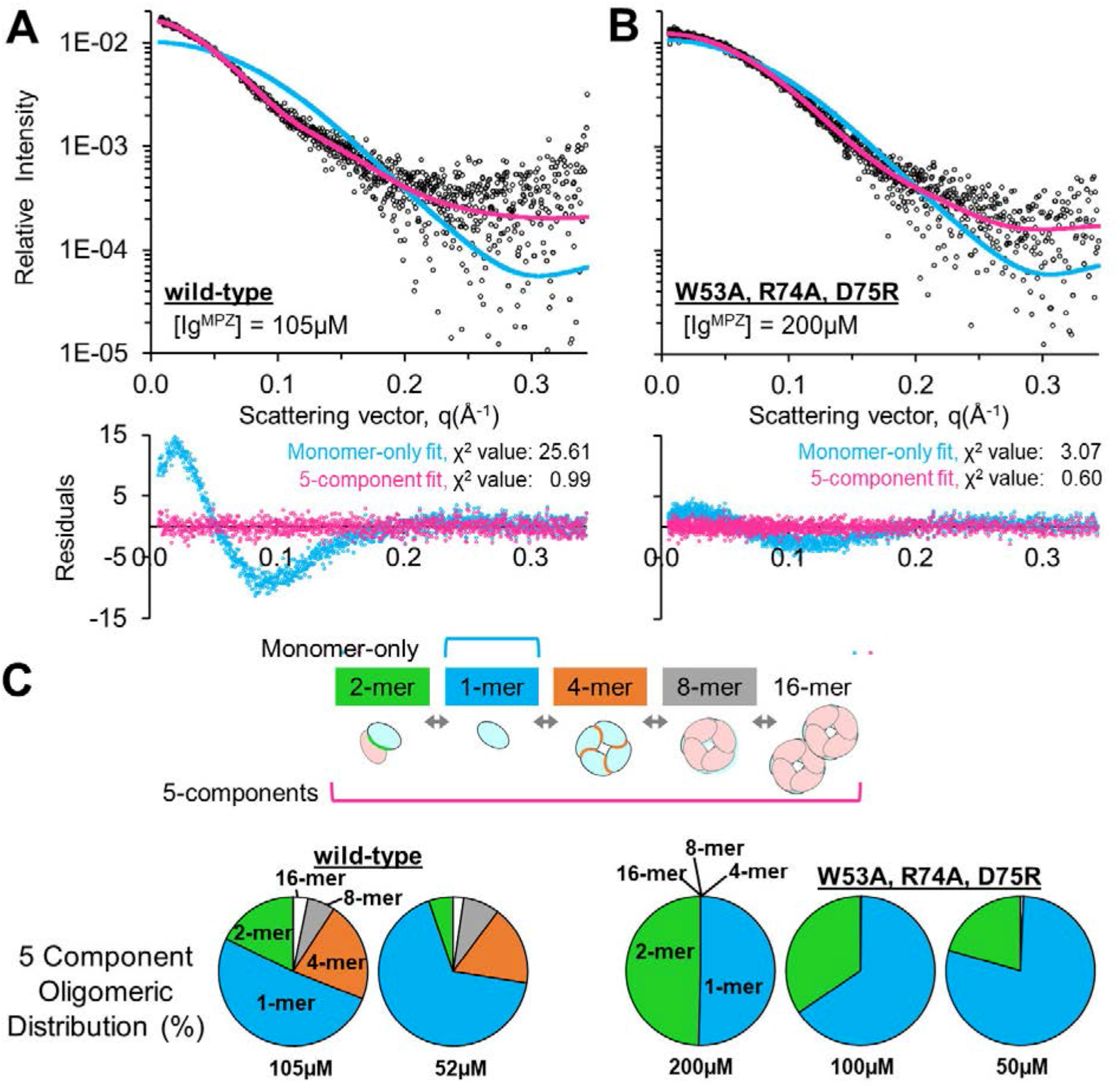
Human lg^MPZ^ oligomeric distribution in solution studied by small angle X-Ray scattering. SAXS data show that wild-type lg^MPZ^ oligomerizes at µM concentrations forming monomers (blue), dimers (green), and tetramers (orange), whereas the lg^MPZ^ ΔAΔB (W53A, R74A, D75R) mutant eliminated tetramer but not dimer formation. (**A**) (Top) SAXS scattering data (circles) for 105 µM wild-type lg^MPZ^. Data are overlayed with the predicted curve for monomeric lg^MPZ^ (blue) and a 5-component fitted curve (pink). (Bottom) Residuals of raw data against monomer only fit (blue) and 5-component curve fit (pink) are shown along with calculated x^2^ values. **(B)** SAXS data (circles), monomer only fit (blue), 5-component curve fit (pink), and residuals for lg^MPZ^ ΔAΔB (W53A, R74A, D75R) measured at 200 µM. **(C)** lg^MPZ^ oligomeric distributions obtained from 5-component fits shown in A and B as well as diluted concentrations of ½x (both) and ¼x (lg^MPZ^ ΔAΔB). Curve fitting components (monomers, dimers, tetramers, dimeric-tetramers, dimeric-octamers) were modeled from the predicted oligomeric arrangement in the crystal structure (PDBid: 1NEU) (Shapiro, et al., 1996). The proportions of each component that best fit experimental data are shown graphically (plotted from data in *Supplemental Table 4*).

We next analyzed the mutant lg^MPZ^flAflB by SAXS (Figure 3B) and confirmed that this mutant still underwent concentration dependent oligomerization as observed in NMR experiments (Figure 2C). However, the extent of lg^MPZ^flAflB oligomerization was less than for wild-type lg^MPZ^, as judged by the difference between experimental data and the theoretical curve of only monomeric lg^MPZ^, and was consistent with the NMR T1/T2 relaxation data which also indicated less oligomerization. The fit of the lg^MPZ^flAflB SAXS data to the theoretical oligomeric distributions optimized for wild-type lg^MPZ^ showed that lg^MPZ^flAflB still formed a dimer in a concentration-dependent manner, but had lost the ability to form tetramers and 8-mers (a dimer of tetramers) (Figure 3C and *Supplemental Table 4*).

Taken together, these data substantiate that lg^MPZ^ forms oligomers in solution and that there are at least 2 interfaces that mediate these interactions. One of the interfaces relies on W53 and supports the idea that the tetrameric lnterface A found in the crystal lattice of the rat lg^MPZ^ in previous studies indeed operates in solution. The data also suggest another interface that mediates lg^MPZ^ dimerization. However, this second interface is not the dimer lnterface B present in the rat lg^MPZ^ crystal lattice (Figure 1A and Figure 2A).

### Distribution of disease-causing variants in the lg domain of MPZ reveals 3 subgroups

Our solution binding data substantiated that the lg^MPZ^ forms tetramers. Moreover, inspection of the transmembrane domain of MPZ shows that it contains two GxxxG motifs known to allow tight packing of transmembrane domains together (Dong et al., 2012; Plotkowski et al., 2007; Senes et al., 2000). Together, these data support the idea that MPZ forms *cis*-tetramers in the membrane. Using a structural bioinformatics approach, we next analyzed the distribution of disease-causing patient amino-acid substitution variants to determine whether there were any distinct subdomains of lg^MPZ^ that correlate with different disease phenotypes or pathogenic mechanisms. Our intent was to determine if a predictive model for types of CMT could be generated based on structural correlates. Moreover, we also wanted to determine whether there were disease-related ‘hot spots’ on the lg^MPZ^ surface that might predict particular protein-protein interaction interfaces such as that which could mediate the unknown ‘dimer’ interface between MPZ tetramers. Previous studies have defined different phenotypes that can arise from variants in MPZ, including just in the lg^MPZ^ domain (Sanmaneechai et al., 2015). One phenotype is classified as CMT2l/J characterized by axonal degeneration but with relatively normal myelin morphology. The other phenotype is classifed as demyelinating CMT1B, some of which can be very early onset and very severe, historically classified as DSS (Dejerine-Sottas Syndrome).

Using patient variant data summarized in Table 1, we calculated two parameters. One was the solvent-accessible surface area as a measure of how close the variants were located to the surface of the protein (Figure 4A). The other parameter was calculated using DeepDDG (Cao et al., 2019) to predict the effect of patient variants on the overall stability of the lg^MPZ^ domain determined by the change in ΔG in kcal/mol (Figure 4B and Table 1). We categorized the patient variants into 2 disease phenotypes, axonal late-onset CMT2l/J or demyelinating early onset CMT1B. We limited analysis to single amino-acid substitutions, excluding deletions, frameshift, or premature stop codon variants that would destroy the lg^MPZ^ domain altogether. We also filtered out variants at positions that can cause different disease phenotypes depending on what residue they are changed to.

**Figure 4.**
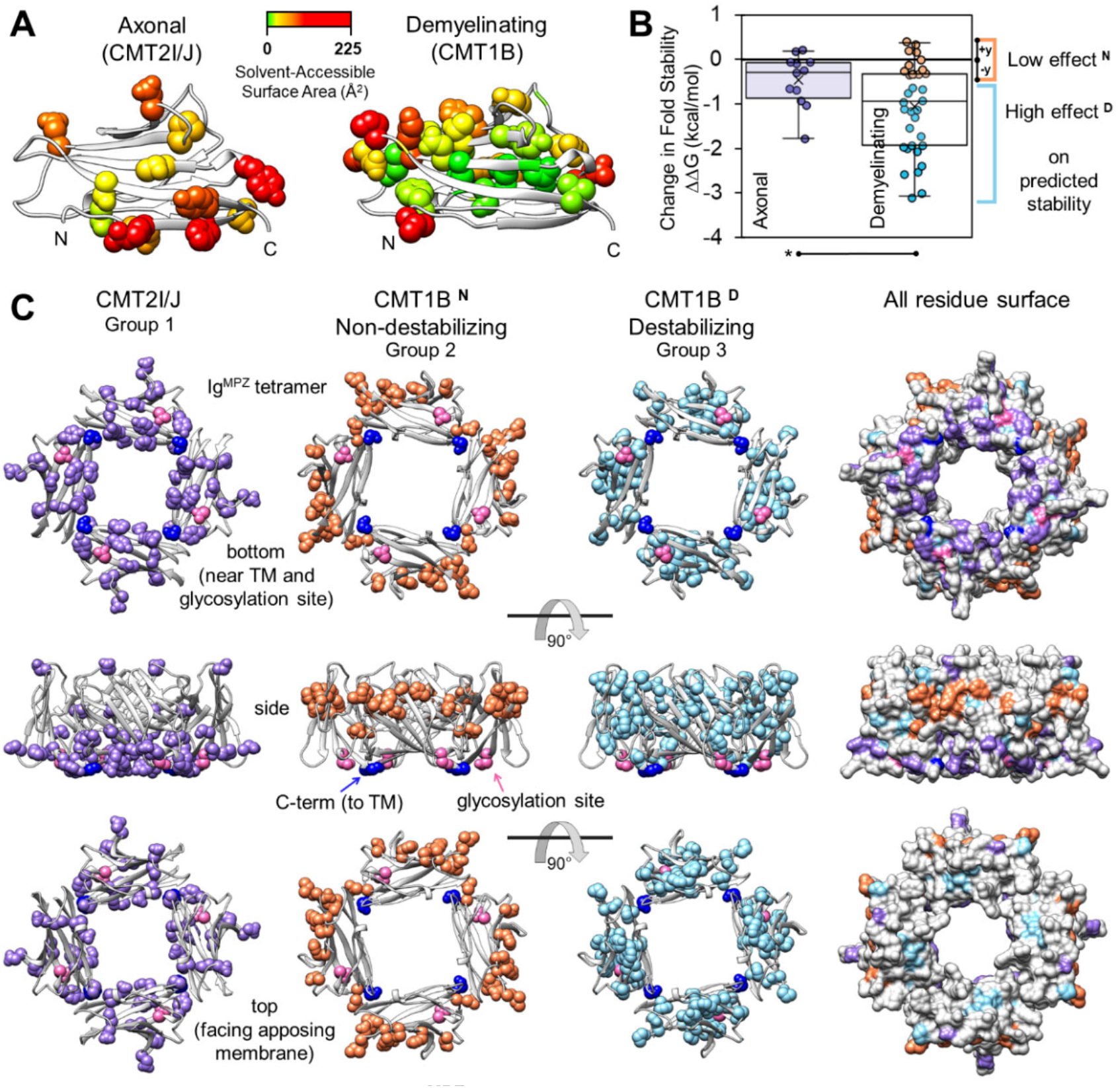
Structurally distinct regions of lg^MPZ^ *cis*-tetramers correlate to disease phenotypes. **(A)** Surface accessibility of wild-type lg^MPZ^ residues with amino acid variants that cause axonal CMT2l/J or demyelinating CMT1B. Positions were mapped onto the human lg^MPZ^ structure (PDBid: 30Al) (Liu, et al., 2012) and color-coded for solvent-accessible surface area. A larger percentage of CMT1B linked residue positions are buried. **(B)** Patient variants causing different CMT phentotypes defined previously (Sanmaneechai et al., 2015) (axonal CMT2l/J or demyelinting CMT1B and listed in Table 1) were computationally evaluated for whether they would destabilize the lg^MPZ^ structure quantified as a change in predicted 11G for lg^MPZ^ stability. Demyelinating CMT1B patient variants had on average a higher prediction for destabilization than axonal CMT2l/J linked variants (*: p<0.05). The CMT1B patient variants could be divided into two subgroups: one with low predicted effects on stability (^N^), and another with high predicted effects on stability (^D^). The surface assessibility of these groups (**(C)** The three groups of residue positions (listed in Table 2) are shown in the context of the *cis* lg^MPZ^ tetramer in 3 viewing orientations. Group 1 (violet) are residues mutated in CMT2l/J; Group 2 (orange) are residues mutated in CMT1B that result in low predicted effects on lg^MPZ^ stability; and Group 3 (light blue) showing positions of CMT1B mutations predicted to destabilize the lg^MPZ^ domain. (Right column) Using the molecular surface of the tetramer, colored patches indicate the solvent accessible residues in the context of surrounding residues. A CMT2l/J patch is visible on the bottom surface of the lg^MPZ^ tetramers near the transmembrane domain and glycosylation site, while a CMT1B^N^ patch is visible along the side view. Surface-inaccessible (buried) residues are not visible in this schematic. *Supplemental Figure 4* shows that the CMT1B^D^ (Group 3) residues have significantly lower levels of surface accssibility than other residue Groups.

**Table 1.**
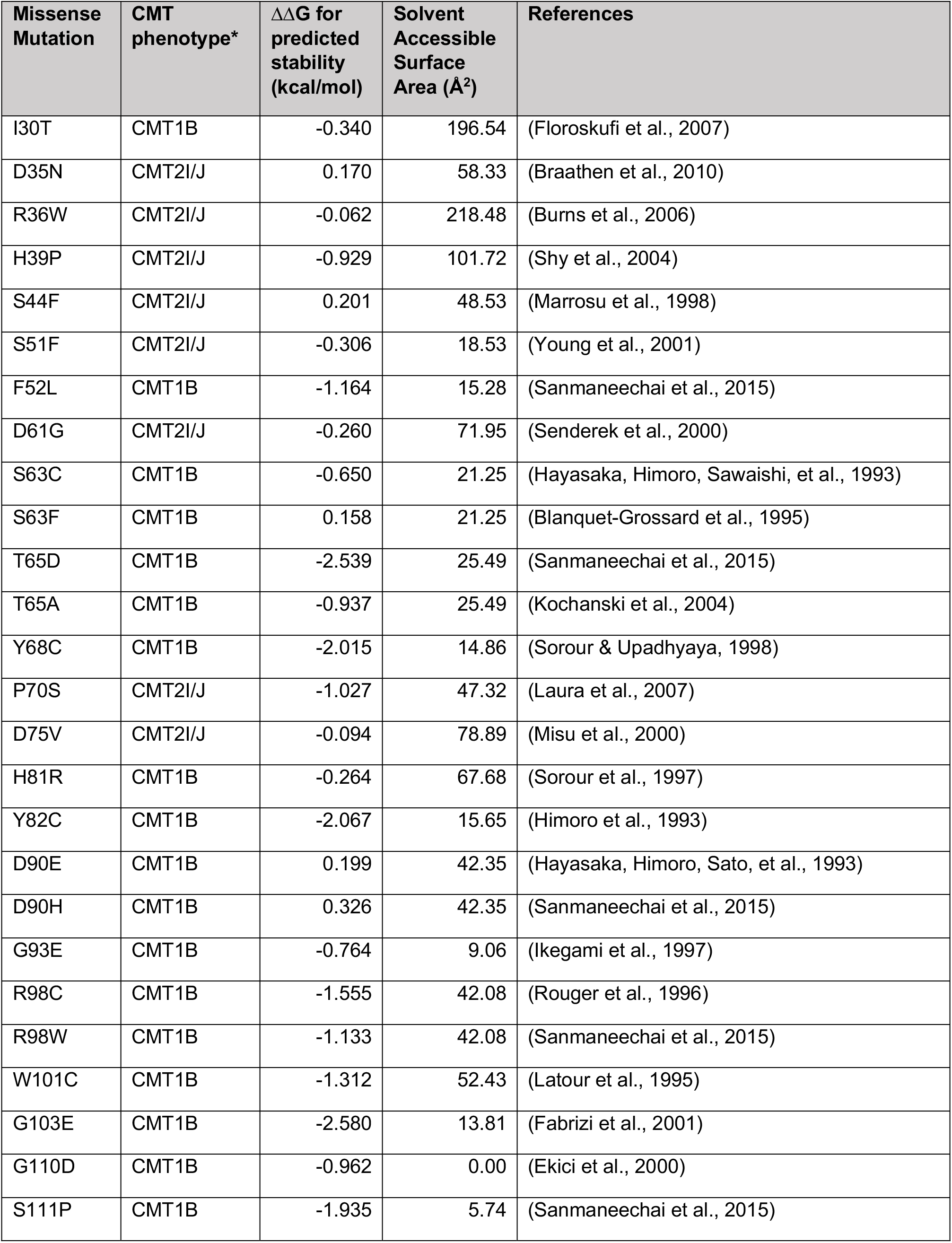
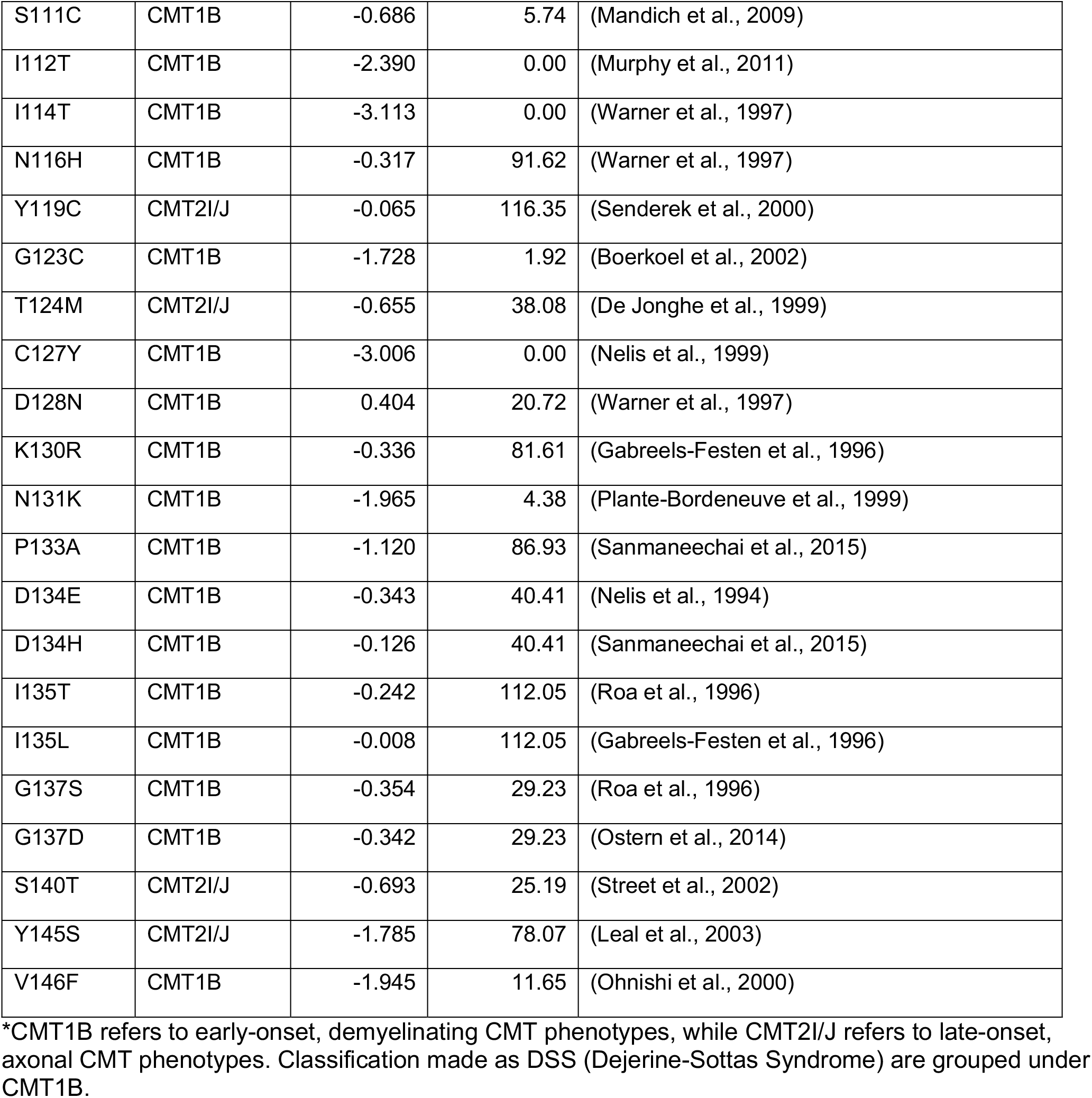
Patient variants evaluated in genotype-structure-phenotype analysis

**Table 2.** Residue position groups defined by CMT phenotype and biophysical parameters

| **Group 1: Residues wherein variants cause axonal CMT2I/J** | **Group 2: Residues wherein variants cause demyelinating CMT1B but NOT predicted to destabilize Ig<sup>MPZ</sup>** | **Group 3: Residues wherein variants cause demyelinating CMT1B and predicted to destabilize Ig<sup>MPZ</sup>** |
| --- | --- | --- |
| D35, R36, H39, S44, S51, D61, P70, D75, Y119, T124, S140, Y145 | I30, S63*, H81, D90, N116, D128, K130, D134, I135, G137 | F52, T65, Y68, Y82, G93, R98, W101, G103, G110, S111, I112, I114, G123, C127, N131, P133, V146 |

\* Average predicted stability for S63 is in Group 2; Includes S63F (Group 2) and S63C (Group 3, near cutoff).

On average, the variants causing CMT2l/J were predicted to have far less of a destabilizing effect on the lg^MPZ^ fold than the CMT1B group of variants, reaching statistical significance of p<0.05 (Figure 4B). Likewise, the positions of the residues involved in CMT2l/J had a higher solvent accessible surface area indicative of their position at the surface of the lg^MPZ^ domain (*Supplemental Figure 4A*), consistent with their lower propensity of surface residues to cause destabilization to a protein fold when mutated (Dill, 1990). We refer to this set of residues as Group 1 (Table 2). Computationally predicting the effect of CMT1B variants on the stability of lg^MPZ^ revealed 2 subgroups. One group had a low predicted effect on lg^MPZ^ stability (non-destablizing CMT1B^N^, which we refer to as Group 2 lg^MPZ^ variant residues). By assuming some inherent variability in variants that did not affect stability, the range of the group with low stability effects was set between the highest positive patient variant value and the identical negative value. Because there is a general correlation between variant stability and surface accessibility for residues in a protein (*Supplemental Figure 4B*), this group also had a high solvent accessible surface area. A separate group contained variants that were predicted to have large destabilizing effects on the lg^MPZ^ fold (destabilizing CMT1B^D^, which we refer to as Group 3 lg^MPZ^ variant residues), had significantly lower levels of solvent accessible surface area (Table 2).

We next mapped these amino acid phenotype groups onto the modeled structure of a *cis*-lg^MPZ^ tetramer to discern if disease phenotypes mapped to discrete subdomains (Figure 4C). Residues (Group 1) that when mutated give rise to CMT2l/J largely map to surface exposed residues that are found clustered on the proximal side nearest to the MPZ transmembrane domain and the membrane surface that tethers lg^MPZ^ domains (Figure 4C and *Supplemental Figure 4*). This surface also surrounds the site where an N-linked glycan of ∼3kDa would be attached. This sizable glycan would likely interact with this surface as well, possibly indicating that the critical nature of this subdomain is to properly engage or accommodate the N-linked glycan.

Group 2 positions were patient variants that cause demyelinating CMT1B but are not predicted to de-stabilize the lg^MPZ^ fold (Table 2). These positions largely map to a distal surface in lg^MPZ^ *cis*-tetramers, away from the transmembrane domain, nearer to the apposing membrane within myelin wraps (Figure 4C). lnterestingly, the ‘*en face*’ view of this subregion shows that it is confined to the outer perimeter of the tetramer surface, ideally suited for interaction with other protein partners. Although this subregion is on the distal end of the lg^MPZ^ tetramer with respect to its position from the membrane, it was distinct from the hydrophobic region at the distal top of the lg^MPZ^ domain that contains hydrophobic tryptophan residues that previous studies proposed may play a role in association with the lipid bilayer of the apposing myelin membrane (Shapiro et al., 1996).

Group 3 positions were sites where patient variants were predicted to destabilize the lg^MPZ^ fold (Table 2). These largely were found in the interior of each lg^MPZ^ subunit within the model tetramer (Figure 4C), consistent with the idea that the amino-acids critical to the integrity of the lg^MPZ^ fold itself would be located within the core of the protein with less surface accessibility.

### Different spatial distribution groups correlate with disease mechanism

One of the best understood pathogenesis pathways for how MPZ mutations can cause dominant-negative CMT is by causing ER-stress and promoting the UPR (Das et al., 2015; Saporta et al., 2012). This occurs when improperly folded proteins accumulate in the ER, up-regulate BiP and trigger signaling through PERK, lRE1and ATF6 (Ron & Walter, 2007). Previous studies in cell culture models have surveyed various patient variants throughout the lg^MPZ^ domain for their ability to elicit UPR (Bai et al., 2018). However, not all CMT1B-causing variants evoke the UPR implying that there are additional mechanisms that can drive demyelinating CMT. The analysis in Figure 4 differentiated a group of CMT1 variants that were predicted to de-stabilize the lg^MPZ^ domain from a group of variants with low predicted affect on the lg^MPZ^ stability. Therefore, we tested whether these computed effects correlated with how these variants evoked the UPR in heterologous expression systems described in previous studies (Bai et al., 2018). To generate a numerical index we combined values that measured UPR transcriptional response as well as the degree of ER localization (*Supplemental Table 5*). We found the destabilizing group of CMT1B variants largely mapping to residues in the core of the lg^MPZ^ (Group 3, CMT1B^O^) had evoked the UPR significantly (p<0.05) more than the non-destabilizing group of CMT1B variants (Group 2, CMT1B^N^) that largely map to surface-exposed residues (Figure 5). These data suggest not only a way to differentiate and predict which lg^MPZ^ variants would trigger a pathogenesis pathway driven by ER-stress, but emphasize that disrupting the function of a different part of the lg^MPZ^, namely the surface on the distal exterior of lg^MPZ^ tetramers (Figure 4), fulfills a distinct function that when compromised leads to a different disease mechanism that drives CMT1B.

**Figure 5.**
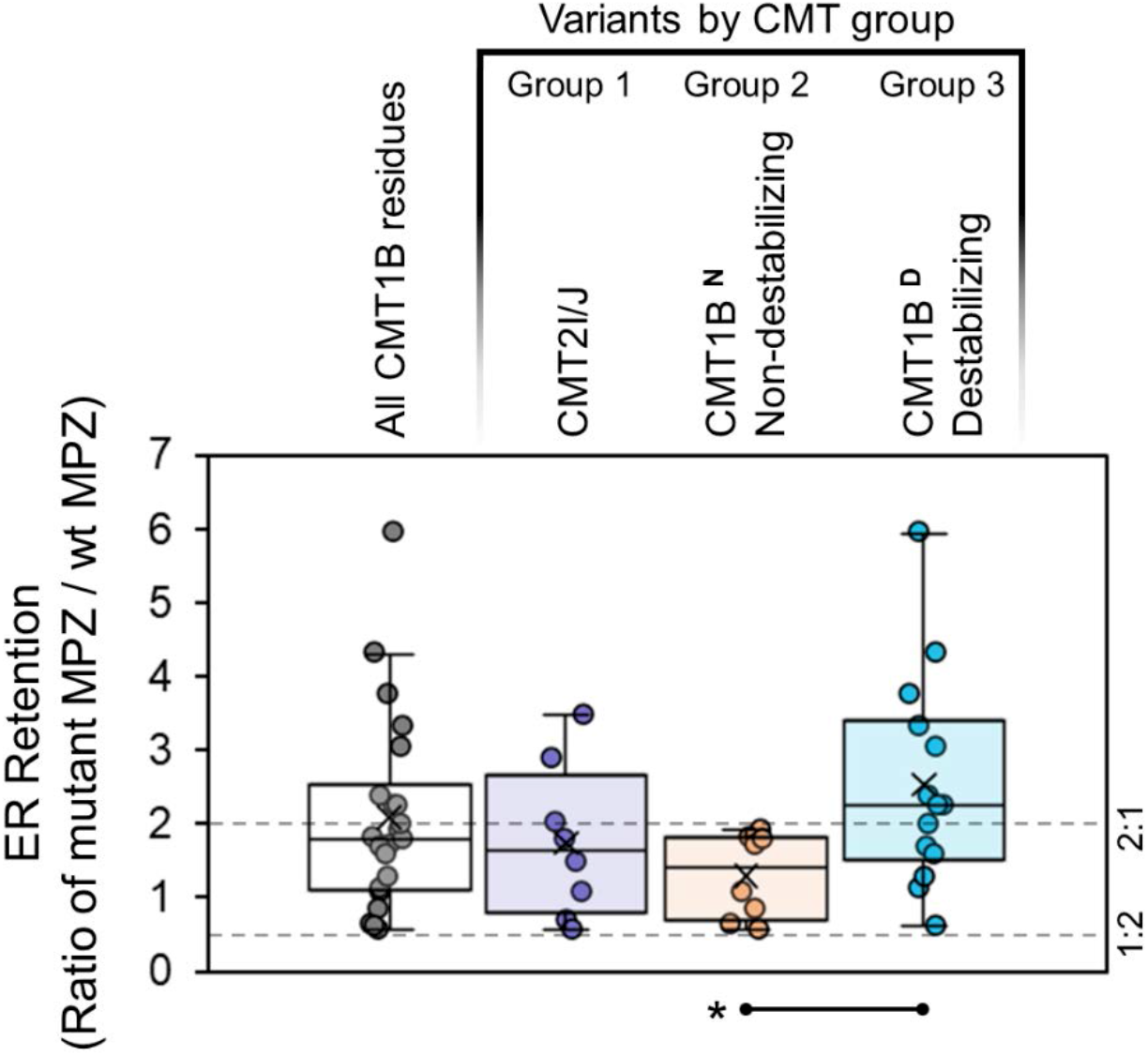
ER retention of MPZ as a function of CMT lg^MPZ^ variant groups. The ability of different CMT-causing MPZ amino acid variants to promote ER retention and activate the Unfolded Protein Response was previously determined (Bai, et al., 2018). For each CMT lg^MPZ^ variant, measurements were averaged and grouped for CMT2l/J (Group 1), and the two CMT1B subgroups (Groups 2 and 3). CMT1B^N^ (Group 2) and CMT1B^O^ (Group 3) were classified as described in Figure 4B and listed in Table 2). The average ER retention and activation of UPR for non-destabilizing and destabilizing CMT1B variant subgroups is significantly different. (*=p<0.05). For variants that did not cause ER retention and UPR had an index in the range between ratio between 2:1 and 1:2 and 2:1 whereas variants that had dramatic ER retention and UPR exceeded 2:1. Data plotted for this evaluation are listed in *Supplemental Table 5*.

### Potential involvement of disease-relevant surface subregions of lg^MPZ^ in dimerization

Analysis in Figure 4 highlighted 2 surface patches on lg^MPZ^ tetramers that when mutated map to 2 different disease phenotypes. Variants in the distal surface cause CMT1B, whereas variants in the proximal surface cause axonal CMT2l/J. Our NMR and SAXS analysis corroborate that lg^MPZ^ *cis*-homotetramerization operates in solution. The identity of a dimer interface that could hold *cis*-tetramers together is distinct from that predicted by previous crystal packing. Therefore, we used computational docking methods to find potential dimer interfaces and then determined whether a particular subregion on the lg^MPZ^ surface was favored in these models. Here, we used ClusPro to calculate a group of 71 dimer poses using one lg^MPZ^ monomer binding to another lg^MPZ^ monomer (Figure 6A)(Kozakov et al., 2017). ClusPro produced 4 sets of poses validated with 4 different scoring schemes that differ in how they weigh energy coefficients (eg favoring electrostatic, hydrophobic, or van der Waals forces). We then filtered these poses to find those that did not produce steric clashes when put into the context of lg^MPZ^ *cis*-tetramers (Figure 6A). This resulted in only 5 tetramer-compatible poses (Figure 6D) in addition to the dimer interface extrapolated from the 1NEU crystal structure (Figure 6C). The 5 new poses were all identified with a scoring scheme that favored a mix of electrostatic interactions combined with van de Waals (shape complementarity). Strikingly, these interfaces involved surface residues from the group (Group 2: CMT1B^N^) of surface-exposed non-destabilizing CMT1B variant group located on the outside distal surface of the lg^MPZ^ *cis*-tetramer (Figure 4C and Table 2). To view this quantitively, we calculated the sum of the distances from all of the 10 CMT1B (Group 2) surface residues to the docked lg^MPZ^ in each dimer pose as well as the same parameter for the 10 CMT2 (Group 1) non-proline surface residues (Figure 6B and *Supplemental Figure 5*). These values convey that the CMT1B^N^ surface (Group 2 residues) was significantly closer to the lg^MPZ^ dimerization interfaces than the CMT2 surface (defined by Group 1 residues) was (*Supplemental Figure 6*). These data also show that 4 of 5 dimer configurations exhibited more engagement with the CMT1B surface than the CMT2 surface as indicated by a lower CMT subgroup proximity score (Figure 6B). Only one pose (6_9 in Figure 6) docked onto the tetramer such that it was closer to the Group 1 ‘CMT2’ surface. However, this pose places two tetramers at an angle that is incompatible with the insertion of the transmembrane domains within either the same or different lipid bilayers across the intraperiod line. This analysis is entirely consistent with the existence of distinct tetramer and dimer interfaces determined by NMR and SAXS providing additional indirect support to a dimer interface that can hold tetramers together exists, but is likely one that is distinct from that modeled from previous crystal packing data. Moreover, the observation that the computational predictions for dimer formation that are compatible within the context of linking tetramers map to surface residues relevant to CMT1B predicts that this surface patch is disease relevant possibly by mediating tetramer-tetramer interactions.

**Figure 6.**
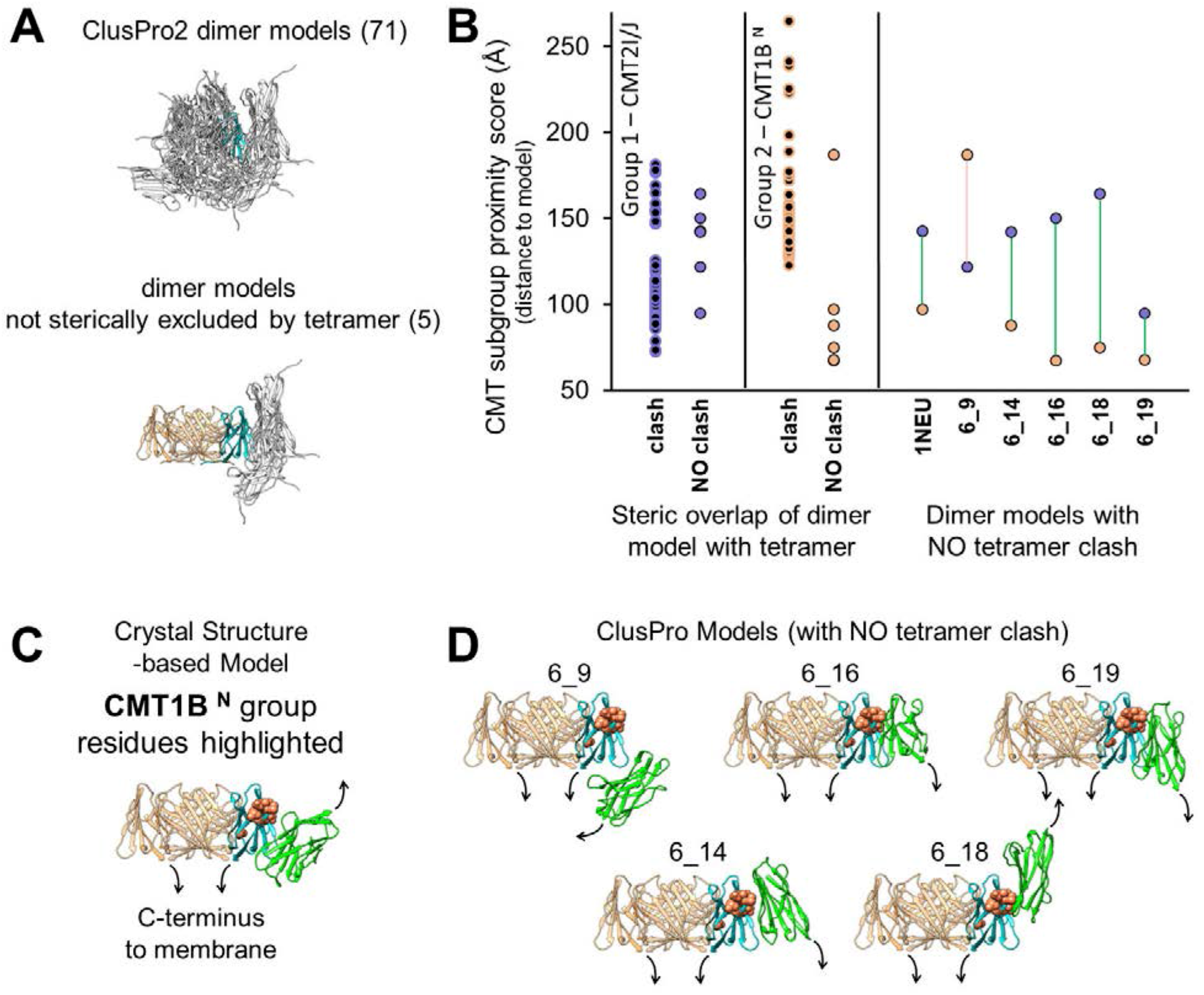
Evaluation of lg^MPZ^ dimer docking models for tetramer compatibility and proximity to CMT subgroup residues. Computational docking experiments reveal that the CMT1B^N^ (Group2) surface subregion may mediate how a cis-lg^MPZ^ tetramer binds (dimerizes) to another tetramer **(A)** Dimer docking poses between lg^MPZ^ monomers calculated using ClusPro (top) were filtered for those that were sterically compatible in the context of lg^MPZ^ tetramers. 5 of 71 dimer arrangements (bottom) could be positioned without overlapping a coexistent *cis* homo-tetramer. **(B)** ClusPro models that had no clash in the context of lg^MPZ^ tetramers or that were excluded because of clash were evaluated for their proximity to CMT2l/J (Group 1) and CMT1B^N^ (Group 2) linked residues. For each model, a subgroup proximity score was calculated by summing the shortest distance between each subgroup residue and the docked lg^MPZ^ for the dimer model being evaluated. (Right) For each of the tetramer compatible dimer models, the subgroup proximity score for CMT2l/J (Group 1) (violet) and CMT1B^N^ (Group 2) (orange) variants were directly compared. **(C)** Cartoon ribbon model for an lg^MPZ^ monomer (cyan) with coexistent *cis* homo-tetramer (orange) and *trans* homo-dimer (green) subunits positioned according to lg^MPZ^ packing in the rat crystal structure (PDBid: 1NEU) (Shapiro, et al., 1996). CMT1B^N^ (Group 2) variant linked residues (orange) are displayed on the cyan lg^MPZ^ subunit. **(D)** The 5 ClusPro lg^MPZ^ dimer docking poses identified in A with the same oligomeric arrangement and color scheme as C.

## DISCUSSION

MPZ is the major membrane protein expressed in Schwann cells and mediates myelin formation. The adhesion activity of MPZ is observed in heterologous cultured cells when ectopically expressed (Filbin et al., 1990; Xu et al., 2001) and MPZ knockout mice have poorly compacted myelin sheaths (Giese et al., 1992). lt is thought that MPZ oligomerizes across the intraperiod line of myelin to hold apposing wraps of Schwann cell plasma membranes together, thus providing a key structural role in forming myelin. Mutations throughout MPZ cause autosomal dominant Charcot-Marie-Tooth disease, a collection of hereditary motor and sensory neuropathies (Fridman et al., 2023; Pisciotta & Shy, 2018; Sanmaneechai et al., 2015; Shy, 2006; Shy et al., 2004), that are characterized either by poorly formed myelin (eg CMT1B) or by axonal degeneration wherein the structure of myelin is relatively intact (CMT2l/J). All of these aspects involve the extracellular lg domain of MPZ (lg^MPZ^) and pose several distinct unanswered questions about how this domain operates in both normal myelinating Schwann cells and in disease pathogenesis.

One key question is to understand the structural basis of lg^MPZ^ oligomerization, its role in normal myelination, and whether perturbation of oligomerization might drive autosomal dominant CMT2 or CMT1B. Current models propose that adhesion of myelin wraps is mediated by lg^MPZ^ oligomerization itself, however this has not been tested (Raasakka & Kursula, 2020; Raasakka et al., 2019; Shapiro et al., 1996). Our data suggest that lg^MPZ^ has two distinct oligomerization modes. One of those is predicted from crystal packing of the rat lg^MPZ^ domain and works by bundling 4 MPZ proteins anchored in the same membrane together, forming a *cis*-tetramer (Figure 1B). Our SAXS data support the model whereby lg^MPZ^ tetramerizes in the ∼100 µM range through an interface that relies on W53 interactions, consistent with the arrangement of tetramers via crystal packing with the rat lg^MPZ^ crystal structure. The lg^MPZ^ with a W53A mutation lost its ability to form tetramers, but retained its ability to oligomerize via dimerization. The interface driving dimerization of lg^MPZ^ domains, however, remains undefined but is distinct from the dimeric interface extrapolated from crystal packing since further targeting this interface did not diminish dimerization. The mode of dimerization which links 2 *cis*-tetramers is important to elucidate since the model that myelin wraps are held together via oligomerized lg^MPZ^ interactions relies on a *trans* interaction of MPZ populating each apposing membrane. While this is a particularly satisfying aspect of both current models for how MPZ mediates adhesion across the intraperiod line, our characterization of lg^MPZ^ in solution requires a different model for how lg^MPZ^ monomers and tetramers would dimerize. Another feature of MPZ noted by Shapiro et al is a hydrophobic patch of Trp residues that might allow the distal surface of an MPZ tetramer to intercalate into the lipid bilayer of the apposing myelin wrap to mediate adhesion (Shapiro et al., 1996). The tetrameric interface that we verify in solution might work to stabilize the orientation lg^MPZ^ domains or increase the avidity of this interface to strengthen this type of interaction. This mode of adhesion is distinct from a *trans* lg^MPZ^-lg^MPZ^ interaction. Our computational studies to find alternate modes of dimerization (Figure 6) only produced one model showing dimerization of tetramers in *trans*, with the rest consistent with *cis*-tetramers linking with other *cis*-tetramers in the same membrane. ln the absence of evidence that the *trans* dimerization interface (lnterface B) mediates oligomerization, alternative models remain distinct possibilities (Figure 7).

**Figure 7.**
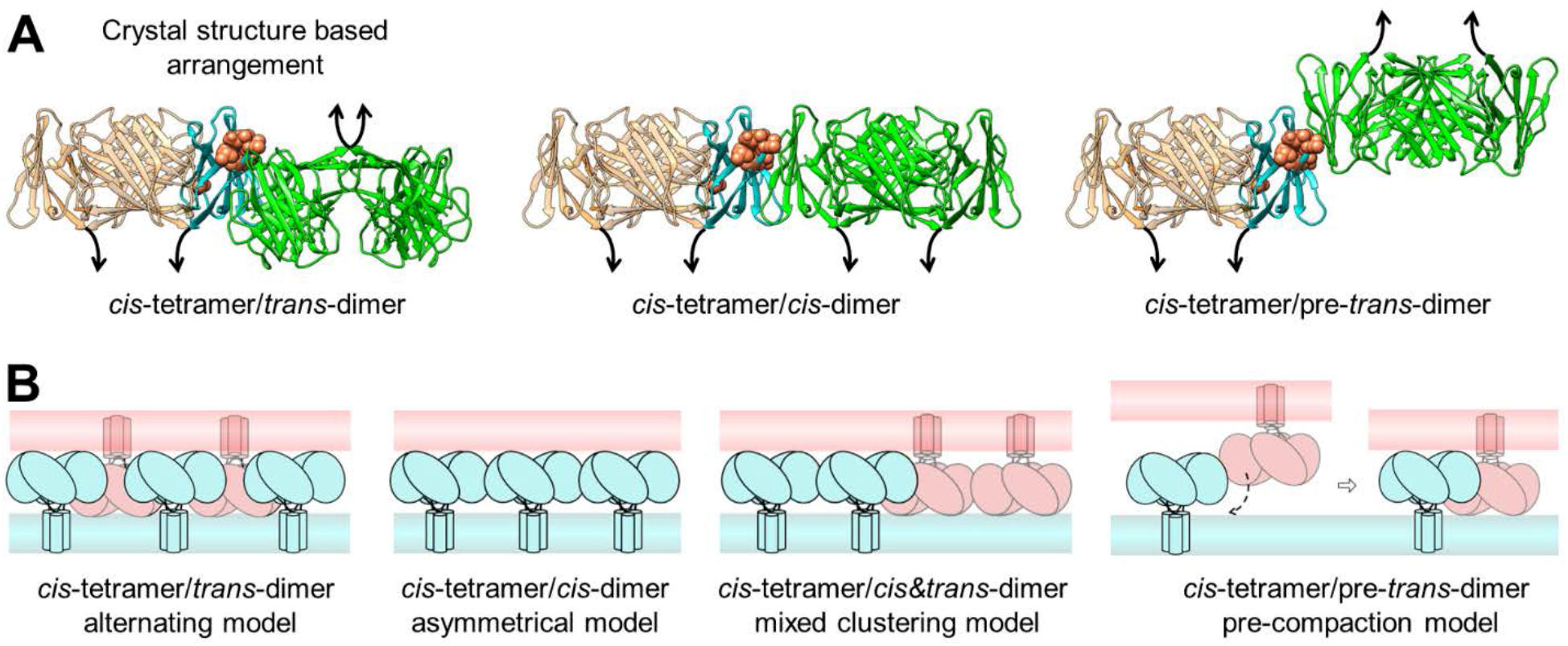
Possible tetramer-tetramer arrangements across intraperiod line. **(A)** Configurations of neighboring lg^MPZ^ homo-tetramers with steric viability across the intraperiod line. (Left) The *cis*-tetramer/*trans*-dimer configuration is based on the packing arrangement in the rat crystal structure. (Middle) An alternate configuration of *cis* interacting *cis*-tetramers, where both lg^MPZ^ homo-tetramers are anchored in the same membrane, is based on the 6_16 ClusPro model in Figure 6D. Models 6_14 and 6_19 suggest similar tetramer-tetramer interaction surfaces. (Right) The 6_18 pose shows the two lg^MPZ^ tetramers in a *trans* configuration but with an interacting surface that is significantly shifted from the crystal structure-based arrangement. The color scheme is the same as in Figure 6. **(B)** Hypothetical schematic arrangements of MPZ within myelin and potential roles of CMT1B^N^ (Group 2) variant linked residues. (Far left) The model developed by Shapiro et al. is based on the lg^MPZ^ crystal contacts. (Middle left) An alternative arrangement incorporates models whereby multiple MPZ tetramers are held together in *cis* via a dimer interface. ln this asymmetrical model, adjacent myelin wraps are stuck together by hydrophobic interactions between the top-distal surface of the lg^MPZ^ tetramer and the lipid bilayer of apposing membrane. (Middle right) A mix of the first two schemes with *trans* interactions between *cis* interacting tetramer clusters is also plausible. (Far right) The final configuration reflects the homo-dimer interaction in the 6_18 ClusPro pose. This tetramer-tetramer interaction would span ∼70 A, which is wider than the ∼45 A span between membrane bilayers across the intraperiod line in compacted myelin and, as such, might represent a transition step during the initial assembly of myelin wraps, to be later reconfigured during myelin compaction.

Recently, Raasakka and Kursula have proposed an alternate model (Figure 1C) for how lg^MPZ^ could mediate adhesion of myelin wraps via zippering of *trans* dimers (Raasakka & Kursula, 2020; Raasakka et al., 2019). That model proposes that the dimer interface is lnterface B but working at a more oblique angle than positioned in the model of Shapiro (Shapiro et al., 1996). Our data support the idea of dimer formation, but not through interface B, and alternative dimer interfaces could not be resolved with the low resolution cryo-EM images Rassakka et al. used to formulate the zipper model (Raasakka et al., 2019). Certainly, other *trans* dimer configurations are possible (e.g., model 6_18, Figure 6), however, the more substantial discrepancy is the omission for a role of lg^MPZ^ tetramers, which clearly form in solution. lt is possible that tetramers might not form in the context of full-length MPZ in biological membranes. Such a conformation requires the transmembrane domains (TMD) of MPZ protomers coming together in close contact, essentially bundling like stems in a flower bouquet of lg^MPZ^ domains. Notably, the membrane-spanning segment of MPZ has two GxxxG motifs that have been found to allow for close packing of transmembrane domains, bolstering the plausibility that *cis* tetramers might form (Dong et al., 2012; Plotkowski et al., 2007; Senes et al., 2000). Another feature of MPZ is its N-linked glycosylation, occurring on a Asn residue within a region of the lg^MPZ^ domain near the membrane that it is inserted into, which would not be predicted to interfere with tetramer formation (Figure 4C). Thus, in our view, the possibility that MPZ forms tetramers in native myelin membranes remains plausible.

Having verified the tetrameric assembly of lg^MPZ^ domains, we also investigated whether there might be functionally distinct regions of lg^MPZ^ that correlate with different disease phenotypes and mechanisms. By computationally calculating which patient missense variants are likely to destabilize the integrity of the lg^MPZ^ fold, we defined 3 discrete subregions of lg^MPZ^ that map to different functions and phenotypes. Variants that cause demyelinating CMT1B generally map to 2 regions. One region was within the core of the lg^MPZ^ and variants there were predicted to have a destabilizing effect on the integrity of the lg^MPZ^ fold (Group 3). Such variants in the core also correlated with their ability to activate the unfolded protein response when expressed in heterologous cells. A different group of CMT1B variants mapped to a surface, lining the exterior distal surface of lg^MPZ^ tetramers where they pose little danger of destabilizing the lg domain but are ideally suited for mediating protein:protein interactions (Group 2: CMT1B^N^). This observation supports the idea of two different disease mechanisms, one being where mis-folded MPZ causes a dominant and deleterious activation of ER-stress. Another distinct mechanism would reflect the function of the CMT1B tetramer surface. Having two distinct CMT1B disease mechanisms mapping to different structural regions of lg^MPZ^ helps rationalize why the severity of CMT1B does not strictly correlate with the extent of UPR activation (Bai et al., 2018). Recently an inhibitor (lFB-088, Sephin1) of the PP1c/PPP1R15A phosphatase complex which works to potentiate UPR has been used to successfully treat mice with CMT1B variants and is entering Phase 2 trials (Bai et al., 2022; Das et al., 2015). We suggest the efficacy of Sephin1 treatment may correlate best with destabilizing variants within the core of the lg^MPZ^ domain vs variants on the surface of the lg^MPZ^ that might cause mis-function via a different mechanism. What the function of the Group 2 ‘CMT1B surface’ is remains to be determined, but it is optimally positioned to mediate protein-protein interaction of an lg^MPZ^ tetramer since these residues line the outside surface of the tetramer. Our computational experiments found that alternate modes of lg^MPZ^ dimerization involved this surface supporting the idea that this region could play a key role in oligomerization and partly explains how perturbation of this surface might contribute to demyelinating disease. Nonetheless, it is less clear whether or how loss of an lg^MPZ^ oligomerization interface might drive autosomal dominant CMT disease. lnability of half of the lg^MPZ^ domains to mediate critical oligomerization events could disrupt the regularity of larger MPZ assemblies that ultimately adhere myelin wraps together, or altering the ability of MPZ to signal to cytosolic components to drive compaction of the major dense line (Previtali et al., 2000; Xu et al., 2001). One possibiity is that trans-assocition could act as an intermediate step, holding the intraperiod line together but with larger spacing until an additional process compresses membrane apposition (Figure 7). lnterestingly, a S63C mutation, located with the CMT1B^O^ surface subregion, appears to cause a disulfide-linkage between trans lg^MPZ^ resulting in a wider 70 A intraperiod line that can be compressed with the disulfide linkage is reduced implying the interface that is locked by a disulfide bond may represent one that is used naturally yet transiently during myelin formation (Avila et al., 2010). Loss of oligomerization might alternatively be recessive, resulting in haploinsufficiency that results in the type of demyelinating and later onset CMT disease that is observed in partial loss-of-function heterozygous MPZ-/+ mice (Martini et al., 1995; Shy et al., 1997) or the more mild neuropathy observed in partial loss-of-function in human CMT patients (Howard et al., 2021). Determining exactly how lg^MPZ^ assembles into oligomers and analyzing the functional consequences of mutations such as W53A that specifically disrupt those interactions will be helpful for resolving these questions.

Spatial mapping also revealed that residues involved in axonal CMT2l/J were on the bottom proximal surface of the lg^MPZ^ tetramer (Group 1). Moreover, patient variants that give rise to CMT2 were not computationally predicted to compromise the stability of the lg^MPZ^ fold, even though some of these variants can evoke the unfolded protein response indicating that the ER quality control system detects a structural problem with them. One key difference here is that while computational studies consider only the lg^MPZ^ protein, when expressed in cells, the lg^MPZ^ domain also undergoes N-linked glycosylation. Moreover, the ‘CMT2-surface’ surrounds the glycosylation site suggesting this region likely interacts with the glycan in a particular way, perhaps to properly position the lg^MPZ^ domain with respect to the membrane. lndeed, mutations in the consensus N-linked glycosylation site lead (N_122_XT) cause late-onset CMT2 phenotypes in humans and mice (Shackleford et al., 2022; Shy et al., 2004) with morphological changes in the areas of non-compacted myelin. Glycosylation has also been found to alter the adhesion activity of MPZ in heterologous systems, suggesting it helps optimally position MPZ for interaction with other components (Filbin & Tennekoon, 1991, 1993). lt is possible that the interaction of the glycan with the CMT2 surface helps position MPZ correctly to control different aspects of Schwann cell development.

## DATA AVAILABILITY

The SAXS experimental data, oligomer fittings, and models from this study are deposited in SASBDB with accession codes: SASDRC3 (lg^MPZ^), SASDRD3 (lg^MPZ^flAflB)

https://www.sasbdb.org/data/SASDRC3

https://www.sasbdb.org/data/SASDRD3

A project summary can be found here: https://www.sasbdb.org/project/1973

## ACKNOWLEDGEMENTS

We thank Dr. Tom Rutkowski (U. Iowa) for helpful discussions and functional insights. We acknowledge the University of Iowa personnel and instrumentation in the IIHG Genomic Sequencing, the Carver College of Medicine NMR, and Protein & Crystallography core facilities, supported by the Roy J. and Lucille A. Carver College of Medicine and grants from the Roy J. Carver Charitable Trust. We acknowledge Sankar Baruah, Gretchen Stennett, and Tabitha Verhage for help with protein purification, Srinivas Chakravarthy, and Maxwell Watkins for help collecting SAXS data, and Manuela Ayee for helpful discussions. This research used resources of the Advanced Photon Source, a U.S. Department of Energy (DOE) Office of Science User Facility operated for the DOE Office of Science by Argonne National Laboratory under Contract No. DE-AC02-06CH11357. BioCAT was supported by grant P30 GM138395 from the National Institute of General Medical Sciences of the National Institutes of Health. The content is solely the responsibility of the authors and does not necessarily reflect the official views of the National Institute of General Medical Sciences or the National Institutes of Health.

This work was supported by NIH-R01 GM106568 to CAA, U54NS065712 to MES, and NIH RO1GM058202 to RCP. CAA and RCP were supported by the Roy J. Carver Charitable Trust.

## AUTHOR CONTRIBUTIONS

CPP: Conceptualization, Methodology, Visualization, Original draft preparation, Investigation.

TAP: Conceptualization, Writing-Original draft preparation, Investigation.

JBH: Conceptualization, Investigation

CAA: Supervision, Conceptualization, Writing-Reviewing and Editing, Funding acquisition.

MES: Supervision, Conceptualization, Writing-Reviewing and Editing, Funding acquisition.

RCP: Supervision, Conceptualization, Writing-Reviewing and Editing, Funding acquisition.

Competing Interests Statement: authors declare no competing interests.

---

**Supplemental Figure 1.**
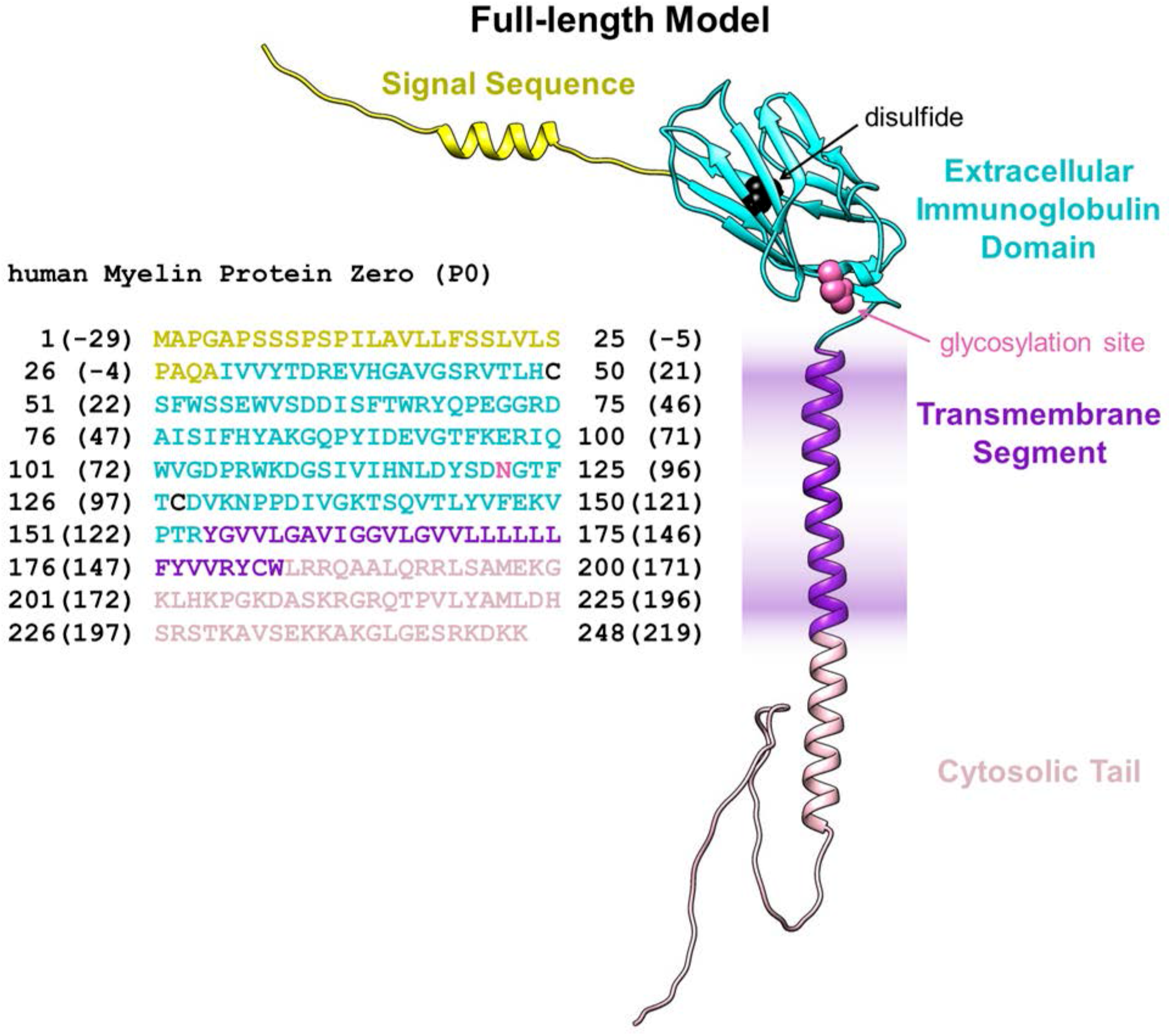
Sequence and organization of human MPZ. Color-coded structural model and sequence of full-length MPZ. Shown is the 29-residue signal sequence (yellow), the 1g domain (cyan), the transmembrane domain segment (purple) and the C-terminal tail (light pink). Residues are displayed for the N-linked glycosylation site (pink) and the internal disulfide bond (black). Along the sequence, residue numbers are shown for the unprocessed MPZ gene product (UniProt P25189) and, in parentheses, the mature MPZ protein after cleavage of the signal-sequence. Note that residue numbers reported for mutations in the literature vary between the full MPZ gene product and the mature cleaved MPZ protein (with 130 as 11). Here, in this manuscript, we refer to the amino-acid numbering scheme of the full gene product including the signal-sequence, which is the current convention.

**Supplemental Figure 2.**
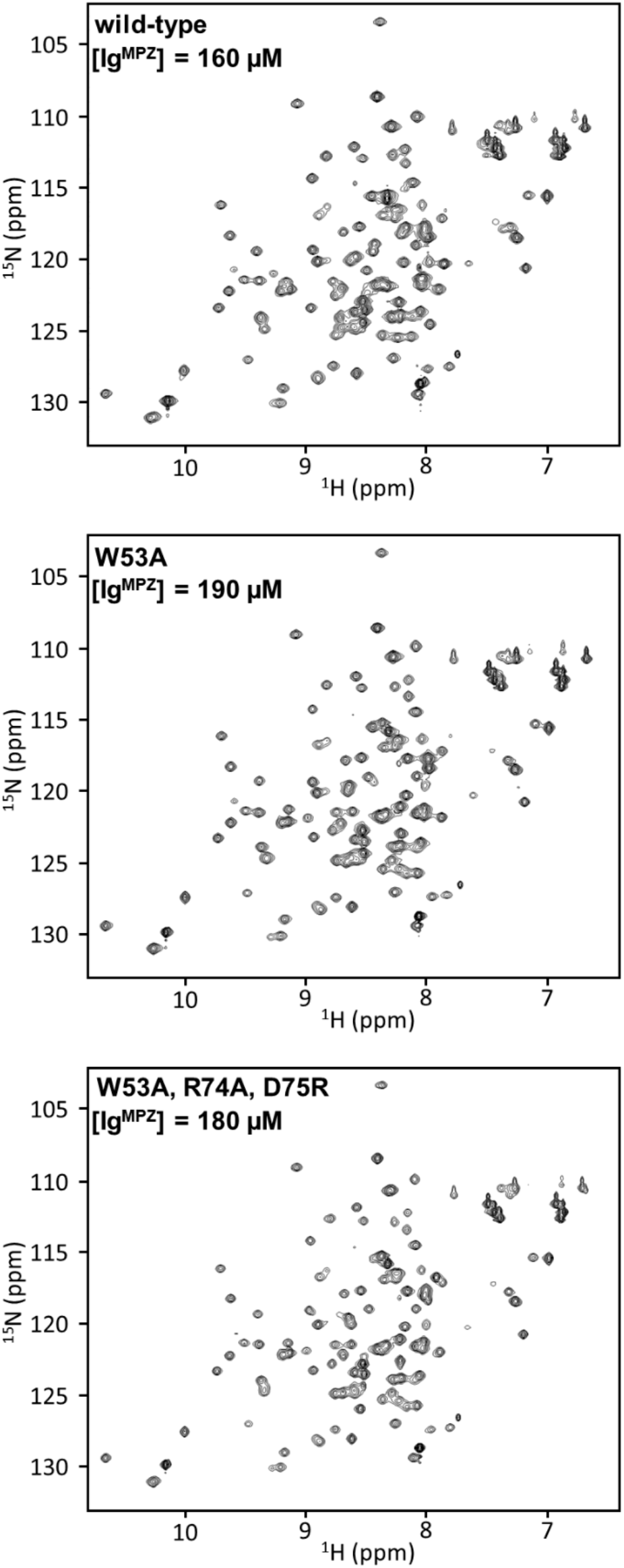
Nuclear magnetic resonance spectra of lg^MPZ^ in solution. Individual ^15^N/^1^H HSQC NMR spectra from the spectral overlay (displayed in Figure 2B) of ^15^N-labeled wild-type lg^MPZ^ (top), the W53A mutant intended to disrupt the Tetrameric Interface A (middle) lg^MPZ^ ΔA, and the combined W53A, R74A, D75R triple mutant targeting both Interface A and the dimer Interface B (bottom) lg^MPZ^ ΔAΔB. Within the set of spectra, the high degree of peak separation and similarity in peak pattern are apparent, while differences between the spectra are easier to observe in the overlay.

**Supplemental Figure 3.**
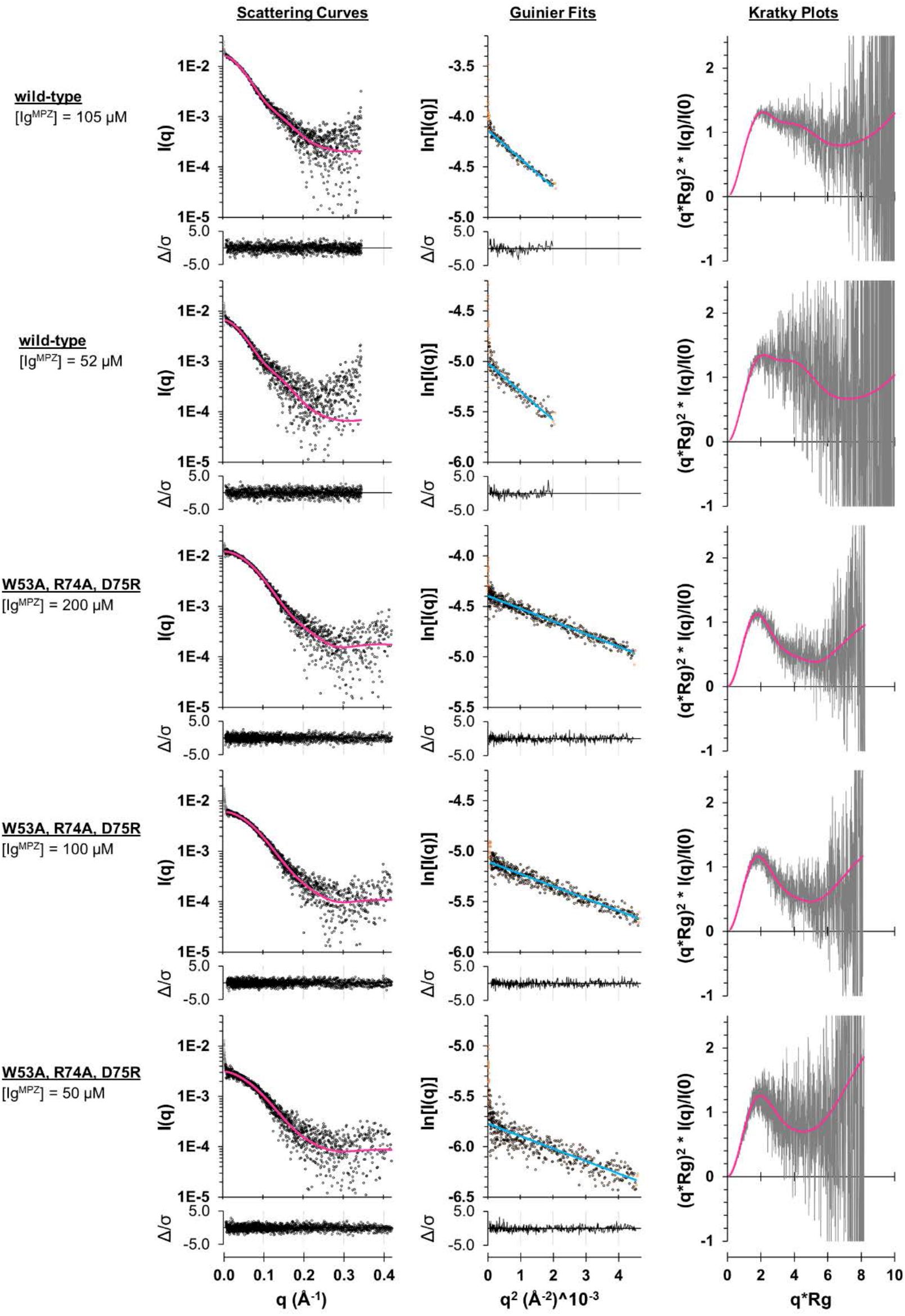
Analysis of SAXS data collected on lg^MPZ^ in solution. For wild-type lg^MPZ^ and the combined W53A, R74A, D75R triple mutant lg^MPZ^ L’AL’B, scattering curves (black circles) are shown (left) with the 5-component fit (pink) and corresponding residuals (below). Guinier analysis (middle) are shown with linear fits (blue) and corresponding residuals (below). The upturn in data at low q is expected for mixtures of molecular species collected in batch mode. The low q data excluded from fitting analysis is included in the scattering curves and Guinier fits (dark gray and orange circles, respectively). The shape of the Kratky plots (right) illustrates the different oligomeric distributions between wild-type lg^MPZ^ and lg^MPZ^ L’AL’B (data - dark gray line; 5-component fit - pink).

**Supplemental Figure 4.**
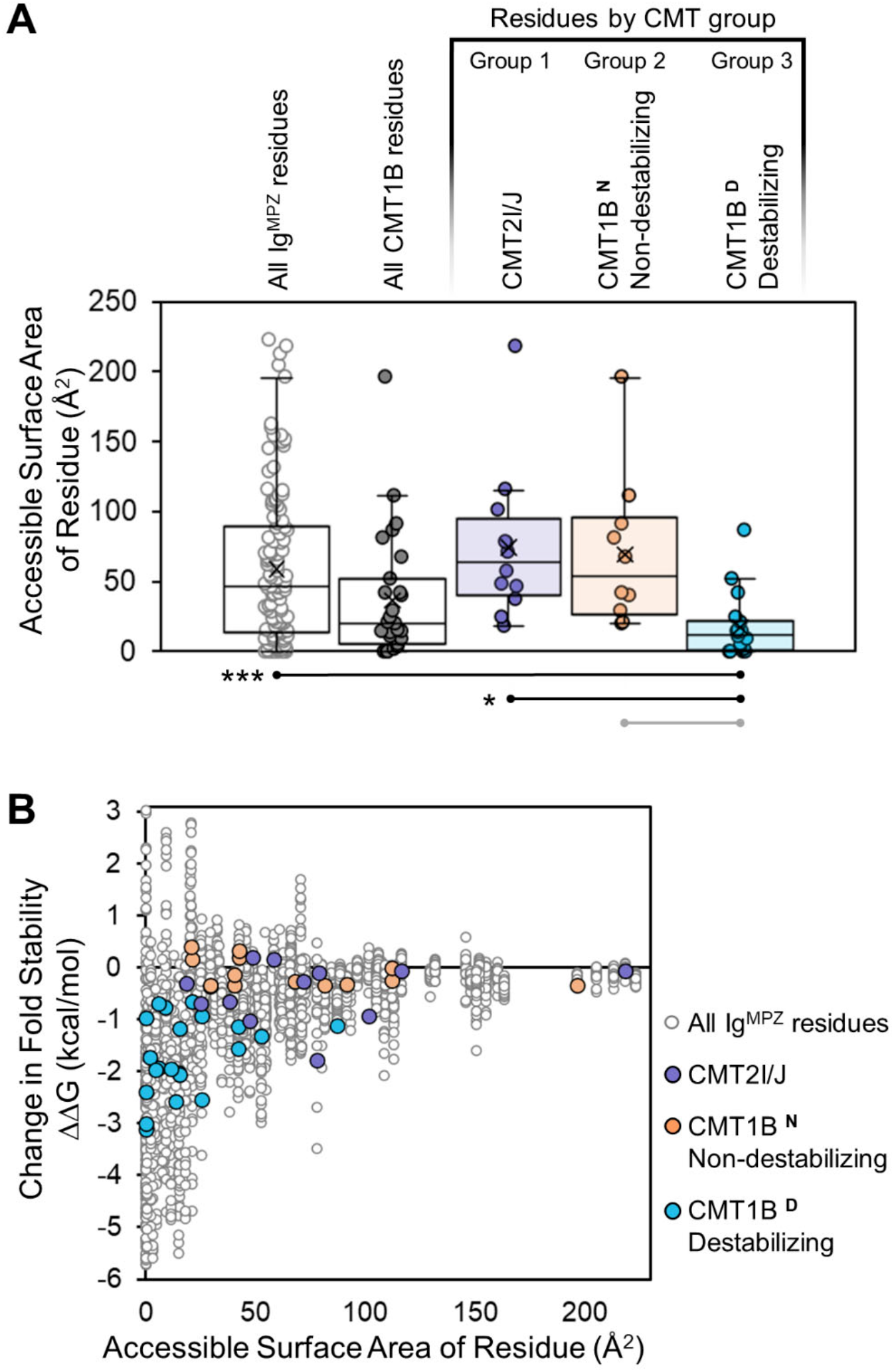
Distribution of predicted destabilizing variants in lg^MPZ^ as a function of solvent accessible surface area. **(A)** Average (±S.D.) of solvent accessible surface area for all the residue positions of lg^MPZ^ (white), those residue positions involved in CMT1B (dark gray), CMT2l/J (violet), and the two subgroups of residues involved in CMT1B, where patient variants had either low (orange) or high (light blue) predicted destabilizing effects on the lg^MPZ^ structure. Group 3, the CMT1B^O^ subgroup with high predicted effect on destabilization, had significantly lower average solvent accessible surface area than the full lg^MPZ^ residue set, Group 1 (CMT2l/J), or Group 2 (CMT1B^N^) (***=p<0.0001; *=p<0.05; grey:p=0.1). **(B)** Plot of change in predicted ΔG for fold stability for all possible variants versus solvent accessible surface area for wild-type residues. The plot shows the expected correlation with variants at buried inaccessible residue positions being more likely to negatively impact protein stability. Data for CMT phenotype subgroup variants are included in the plot.

**Supplemental Figure 5.**
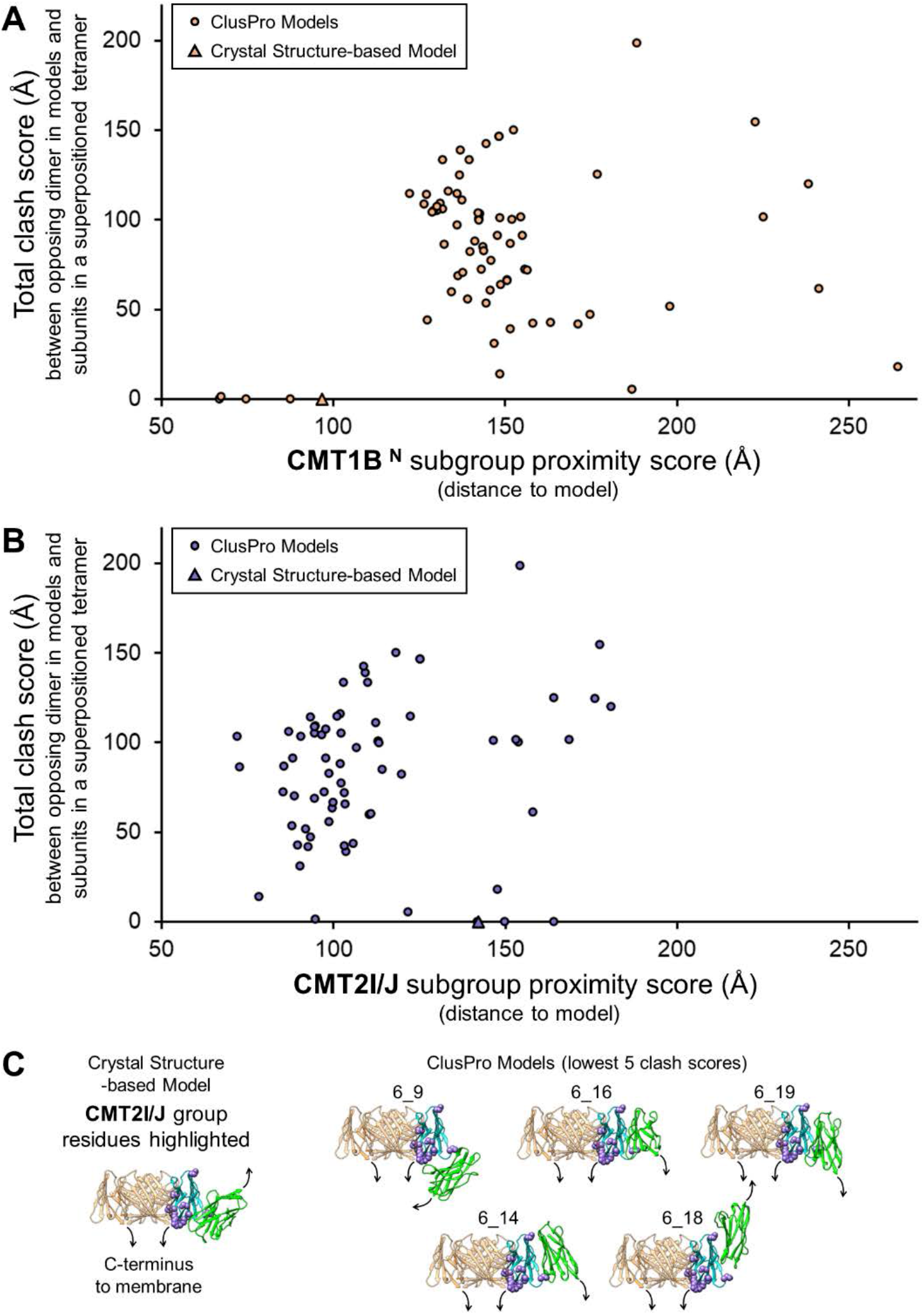
Distribution of tetramer overlap for lg^MPZ^ dimer models relative to model distance to CMT subgroup residues. **(A)** lg^MPZ^ tetramer clash for ClusPro models versus model distance to phenotype subgroup residue sets for CMT1B^N^. The total clash score was calculated for all ClusPro dimer models in the context of a coexistent lg^MPZ^ tetramer. A very low to zero clash score is necessary for the dimer model to be compatible with a tetramer-tetramer interaction site. For each model, a subgroup proximity score was calculated by summing the shortest distance between each CMT1B^N^ (Group 2) residue and the docked lg^MPZ^ for the dimer model being evaluated. The same plot for model distance to the CMT2l/J Group 2 with10-residues (prolines not included) is shown in **(B)**. **(C)** Cartoon ribbon model for an lg^MPZ^ monomer (cyan) with coexistent *cis* homo-tetramer (orange) and *trans* homo-dimer (green) subunits positioned according to lg^MPZ^ packing in the rat crystal structure (PDBid: 1NEU) (Shapiro, et al., 1996). CMT2l/J (Group 1) variant linked residues (violet) are displayed on the cyan lg^MPZ^ subunit. The 5 ClusPro lg^MPZ^ dimer docking poses with no tetramer clash are also shown.

**Supplemental Figure 6.**
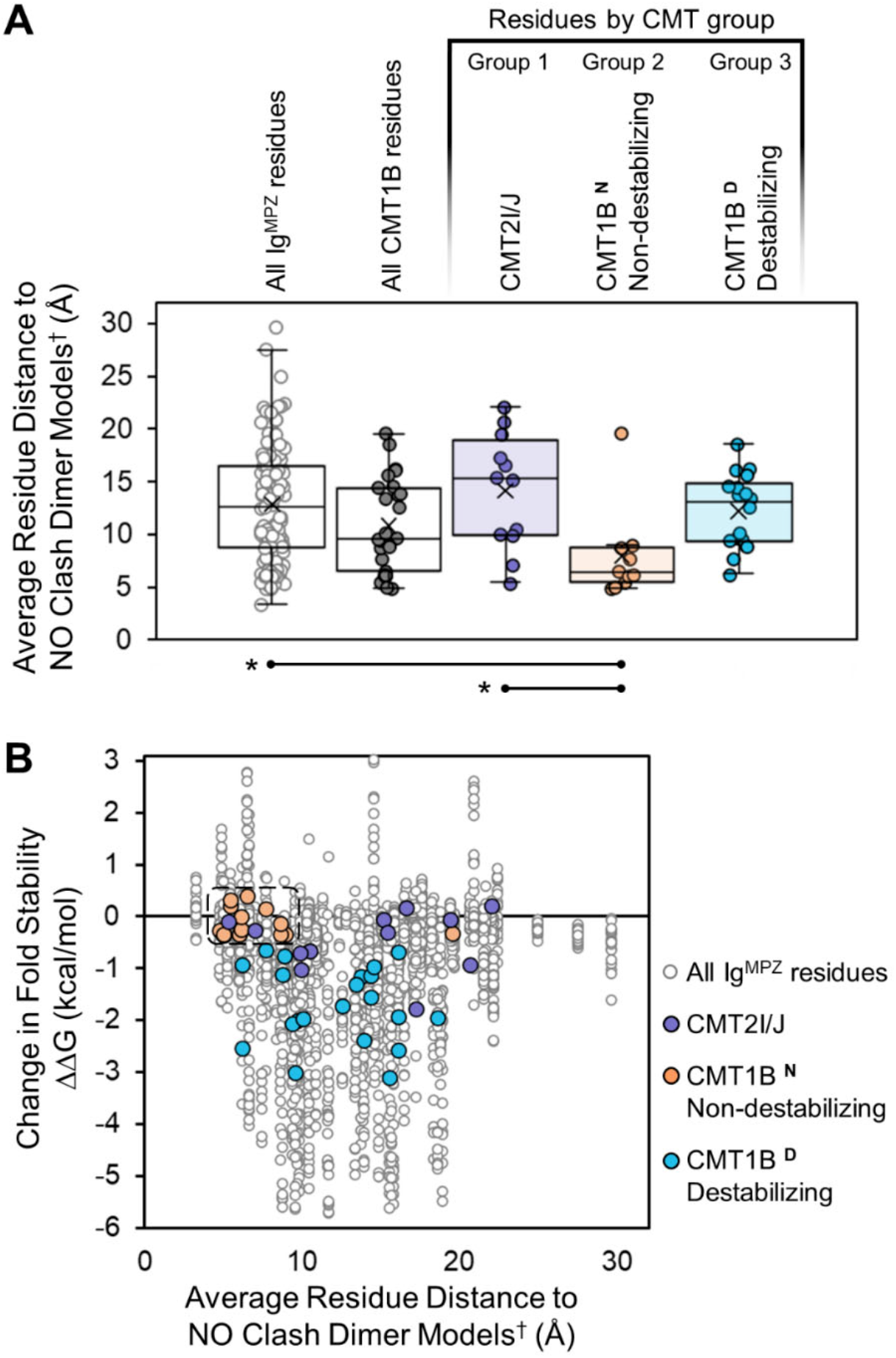
Distribution of lg^MPZ^ residues as a function of distance to NO clash models. **(A)** Averaged distance between lg^MPZ^ residues and each practical models with no tetramer clash (model set: 1NEU-based, 6_14, 6_16, 6_18, 6_19) for all the residue positions of lg^MPZ^ (white), those residue positions involved in CMT1B (drak gray), CMT2l/J (violet), and the two subgroups of residues involved in CMT1B, where patient variants had either low (orange) or high (light blue) predicted destabilizing effects on the lg^MPZ^ structure. Group 2, the CMT1B^N^ subgroup with low predicted effect on destabilization, residues were on average significantly closer to models compatible with interacting tetramers than the full lg^MPZ^ residue set or Group 1 (CMT2l/J) (*=p<0.05). **(B)** Plot of change in predicted ΔG for fold stability for all possible variants versus subgroup residues to dimer model distance. The plot shows no correlation between predicted effect on stability and average residue distance to dimer models with no tetramer clash. Data for CMT phenotype subgroup variants are included in the plot. CMT1B^N^ subgroup variants are almost exclusively part of a tight data cluster suggestive of a patch of surface residues.

**Supplemental Table 1.** Plasmids used in this study.

| Plasmid | Marker | Description | Reference/Source |
| --- | --- | --- | --- |
| pPL7238 | Kan | Secreted-MBP-hMPZ Ig for production in Origami 2(DE3) bacteria | This Study |
| pPL7370 | Kan | Secreted-MBP-hMPZ Ig W53A for production in Origami 2(DE3) bacteria | This Study |
| pPL7481 | Kan | Secreted-MBP-hMPZ Ig W53A, D74A, D75R for production in Origami 2(DE3) bacteria | This Study |
| pPL6973 | Amp | Synthetic DNA encoding human MPZ | This Study |
| pPL7430 | Amp | MPZ-eGFP-HA expressed from CMV promoter | This Study |
| pPL7516 | Amp | MPZ W53A, R74A, D75R-eGFP-HA expressed from CMV promoter | This Study |

**Supplemental Table 2.** Predicted changes in binding energy upon mutation. Computed change in binding energy predicted for different mutations in lg^MPZ^ for the tetramer binding interface (lnterface A) and the dimer interface (lnterface B) found within the crystal packing of the rat lg^MPZ^ structure (PDBid: 1NEU). lnterface B mediates homotypic dimerization where the residues on one lg^MPZ^ subunit mediate binding through the same residue positions on the other lg^MPZ^ binding partner. Thus, the table calculates the change in binding energy (1111G) when either one or both dimerizing (lnterface B) lg^MPZ^ are mutant. lnterface A is a heterotypic binding interface with Y33 and W53 of one lg^MPZ^ subunit interacting with H115, N116, E97 of another lg^MPZ^. Here, we predicted the 1111G of mutating the surface of just one of the two lg^MPZ^ domains at this interface.

| Computed mutations | Computed impact on binding<br>$\Delta\Delta G$ (kcal/mol) | | |
| --- | --- | --- | --- |
|  | Interface A | Interface B<br>1 of 2 chains | Interface B<br>both chains |
| Y33A | 2.154 | 0 | 0 |
| **W53A ($\Delta A$)** | 3.139 | 0 | 0 |
| E97A | 0.511 | 0 | 0 |
| H115A | 1.567 | 0 | 0 |
| N116A | 0.66 | 0 | 0 |
| R67A | 0 | 0.501 | 0.797 |
| Q69A | 0 | 0.544 | 0.883 |
| R74A | 0 | 0.682 | 1.158 |
| D75R | 0 | 0.466 | 1.048 |
| D75A | 0 | 0.205 | 0.865 |
| S78A | 0 | 0.619 | 1.037 |
| **R74A, D75R ($\Delta B$)** | 0 | 0.943 | 2.001 |
| **W53A, R74A, D75R**<br>**($\Delta A\Delta B$)** | 3.139 | 0.943 | 2.001 |
| **Patient variants** |  |  |  |
| N116H | 0.565 | 0 | 0 |

**Supplemental Table 3.**
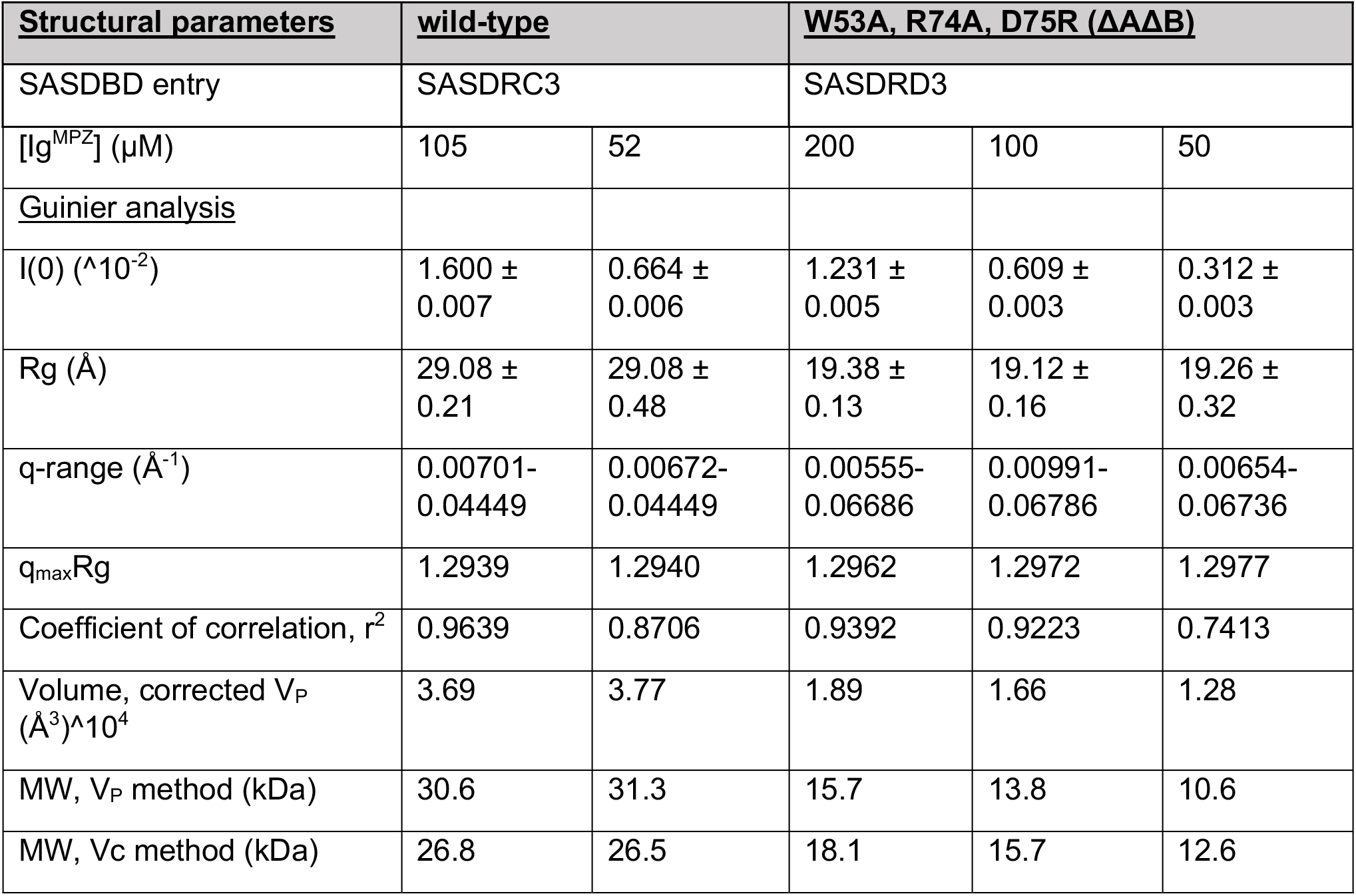
SAXS data analysis parameters.

**Supplemental Table 4.** Proportion of lg^MPZ^ monomeric and oligomeric forms that best fit experimental SAXS data.

| **[Ig<sup>MPZ</sup> wt]** | **<u>2-mer</u>** | **<u>1-mer</u>** | **<u>4-mer</u>** | **<u>8-mer</u>** | **<u>16-mer</u>** | **<u>χ<sup>2</sup></u>** |
| --- | --- | --- | --- | --- | --- | --- |
| 52 μM | 5.3% | 67.2% | 17.3% | 7.9% | 2.3% | 0.995 |
| 105 μM | 17.8% | 51.2% | 21.8% | 6.0% | 3.2% | 0.993 |
| **<u>[Ig<sup>MPZ</sup> ΔAΔB]</u>**<br>**<u>W53A, R74A, D75R</u>** |  |  |  |  |  |  |
| 50 μM | 20.6% | 78.7% | 0% | 0% | 0.7% | 0.557 |
| 100 μM | 34.5% | 65.3% | 0% | 0% | 0.2% | 0.565 |
| 200 μM | 49.6% | 50.4% | 0% | 0% | 0% | 0.599 |

**Supplemental Table 5.** Indices for ER retention and UPR activation calculated from previous data. An index was made from pervious data (Bai et al., 2018) that examined disease variants in MPZ and measured the level of ER-retention and the level of UPR activation. Variant to wild-type ratios from these assays were averaged to minimize assay specific artifacts.

| Missense Mutation | CMT phenotype | Firefly1* | Nano2* | IHC-ER3* | ER retention Index |
| --- | --- | --- | --- | --- | --- |
| R36W | CMT2I/J | 2.7 | 3.5 | 2.5 | 2.9 |
| H39P | CMT2I/J | 1.5 | 2 | 1.9 | 1.8 |
| S44F | CMT2I/J | 1.6 | 1.7 | 1.2 | 1.5 |
| F52L | CMT1B<sup>D</sup> | 1.5 | 1.9 | 1.7 | 1.7 |
| S63F | CMT1B<sup>N</sup> | 1.8 | 2.3 | 1.7 | 1.9 |
| T65N | CMT1B<sup>D</sup> | 1 | 1.3 | 1.1 | 1.1 |
| T65A | CMT1B<sup>D</sup> | 4.6 | 5.1 | 3.3 | 4.3 |
| P70S | CMT2I/J | 1.6 | 1.8 | 2.7 | 2.0 |
| Y82C | CMT1B<sup>D</sup> | 1.1 | 1.4 | 2.3 | 1.6 |
| D90H | CMT1B<sup>N</sup> | 1.5 | 2 | 2 | 1.8 |
| R98C | CMT1B<sup>D</sup> | 1.9 | 2.2 | 2.7 | 2.3 |
| R98W | CMT1B<sup>D</sup> | 5.7 | 7.1 | 5.1 | 6.0 |
| G103E | CMT1B<sup>D</sup> | 3.1 | 4.2 | 2.7 | 3.3 |
| G110D | CMT1B<sup>D</sup> | 3.3 | 4.1 | 3.9 | 3.8 |
| S111P | CMT1B<sup>D</sup> | 2.7 | 3.6 | 2.9 | 3.1 |
| S111C | CMT1B<sup>D</sup> | 1.9 | 2.7 | 2.6 | 2.4 |
| I112T | CMT1B<sup>D</sup> | 1.2 | 1.6 | 1.1 | 1.3 |
| I114T | CMT1B<sup>D</sup> | 1.8 | 2.3 | 1.9 | 2.0 |
| Y119C | CMT2I/J | 0.6 | 0.8 | 0.7 | 0.7 |
| G123C | CMT1B<sup>D</sup> | 2.2 | 2.9 | 1.7 | 2.3 |
| T124M | CMT2I/J | 0.9 | 1.1 | 1.3 | 1.1 |
| K130R | CMT1B<sup>N</sup> | 1.6 | 2.2 | 1.4 | 1.7 |
| P133A | CMT1B<sup>D</sup> | 0.5 | 0.6 | 0.8 | 0.6 |
| D134E | CMT1B<sup>N</sup> | 1.1 | 1.3 | 0.9 | 1.1 |
| D134H | CMT1B<sup>N</sup> | 1.5 | 1.6 | 2.3 | 1.8 |
| I135T | CMT1B<sup>N</sup> | 0.5 | 0.7 | 0.8 | 0.7 |
| G137S | CMT1B<sup>N</sup> | 0.8 | 1.1 | 0.7 | 0.9 |
| G137D | CMT1B<sup>N</sup> | 0.4 | 0.6 | 0.7 | 0.6 |
| S140T | CMT2I/J | 0.4 | 0.6 | 0.7 | 0.6 |
| Y145S | CMT2I/J | 3.5 | 3.7 | 3.3 | 3.5 |

\*Previously determined values (Bai, et al., 2018).

